# Mechanisms of memory storage and retrieval in hippocampal area CA3

**DOI:** 10.1101/2023.05.30.542781

**Authors:** Yiding Li, John J. Briguglio, Sandro Romani, Jeffrey C. Magee

## Abstract

Hippocampal area CA3 is thought to play a central role in memory formation and retrieval. Although various network mechanisms have been hypothesized to mediate these computations, direct evidence is lacking. Using intracellular membrane potential recordings from CA3 neurons and optogenetic manipulations in behaving mice we found that place field activity is produced by a symmetric form of Behavioral Timescale Synaptic Plasticity (BTSP) at recurrent synaptic connections among CA3 principal neurons but not at synapses from the dentate gyrus (DG). Additional manipulations revealed that excitatory input from the entorhinal cortex (EC) but not DG was required to update place cell activity based on the animal’s movement. These data were captured by a computational model that used BTSP and an external updating input to produce attractor dynamics under online learning conditions. Additional theoretical results demonstrate the enhanced memory storage capacity of such networks, particularly in the face of correlated input patterns. The evidence sheds light on the cellular and circuit mechanisms of learning and memory formation in the hippocampus.

**One Sentence Summary:** Evidence from behaving mice points to cellular and circuit mechanisms that underlie observed attractor dynamics in area CA3.

## Introduction

The mammalian hippocampus plays a crucial role in episodic memory formation and retrieval^1-4^. One subregion, area CA3, is thought to be particularly important in this process as consistent, environmentally specific sequences of robust place cell activity originate here^5,6^. The concept of attractors, which are the minimal set of states in a state space to which all nearby states eventually flow, currently represents an appealing theoretical mechanism for memory storage within brains^7-8^. Attractor networks produce a complete output from only a partial set of inputs (pattern completion), have the potential to robustly store and accurately retrieve a very large number of activity patterns, and have been implicated in the creation of distinct output patterns even when presented with similar inputs (pattern separation)^7-9^. While various forms of attractor dynamics have been hypothesized to underlie the mnemonic functions of the hippocampus, whether and how they are implemented in CA3 remains unknown^6,8-17^.

Most relevant to hippocampal memory are networks whose activity dynamics is linked to or updated by the animal’s behavior (fig. S1, A to F). Formation of place-related activity patterns in such networks requires specific adjustments of synaptic strengths to produce a set of neurons that become active together (an activity “bump”) (fig. S1B) and a separate mechanism to associate this neuronal activity with certain internal (movement velocity and path integration) and external (sensory environmental elements) features such that the dynamics of the activity bump is linked to behavior^8-16^ (fig. S1, C to E). This updating or linking mechanism is frequently an additional appropriately tuned excitatory input. The specific adjustments mentioned above are mediated by learning rules that yield synaptic weight changes, usually at recurrent connections, that are symmetrical in network space^8-19^ (fig. S1, A to F). Such rules produce stable network activity dynamics that can give rise to single neuron place-specific activity that persists within the same location in the absence of the additional updating mechanism. Asymmetric learning rules, on the other hand, cause an unstable network dynamics where activity can change independent of any external linking inputs and is thus untethered to behavior^20-21^(fig. S1, G to K). While there are many theories about what learning rules and which pathways allow CA3 to produce the observed network dynamics^6-19,23,24^, there is no relevant direct *in vivo* experimental evidence. We therefore sought to determine 1) what plasticity forms, 2) at which synapses are responsible for individual CA3 place field activity, 3) what input pathways, if any, act as an activity update mechanism and 4) what are the theoretical capabilities of such an attractor network.

### Synaptic mechanisms of CA3 place fields (PFs)

We began by using whole-cell intracellular membrane potential (Vm) recordings from a set of CA3 pyramidal neurons (n=185) in head-fixed mice running laps for a water reward on a treadmill (∼180 cm in length) (Fig. 1A and fig. S2, A and B)^25-28^. Approximately one quarter of the principal neurons (46/185) showed spatially localized action potential (AP) firing during running (Fig. 1B and fig. S2 C to H). This spatially localized firing was associated with a ramping of Vm depolarization (Vm ramp; fig. S2D) and intracellular theta frequency oscillation amplitude (Vm theta; fig. S2E) that both peaked around the same spatial location as the AP rate. In addition to standard AP firing, high frequency burst firing associated with moderate duration after-depolarizations (ADPs) was observed within the PF (fig. S2F). Population averages of AP rate, Vm ramp and Vm theta were generally flat across the track (Fig. 1, C to E) except for a modulation of AP rate perhaps by animal running speed (R^2^=0.81; p<5.4e-36; fig. S2G). These data suggest that the spatial density profile for these CA3 place cells (PCs) was uniform across the environment and this was confirmed by a histogram of PF peak location (Fig. 1F). Together the above results indicate that, like CA1 PCs, PF firing in CA3 is driven by a slow ramp of Vm depolarization and an increase in the amplitude of theta frequency Vm oscillations. However, unlike CA1 the spatial density of CA3 PCs appears to be uniform across the environment, which is a favorable condition for stable attractor dynamics.

**Fig. 1.**
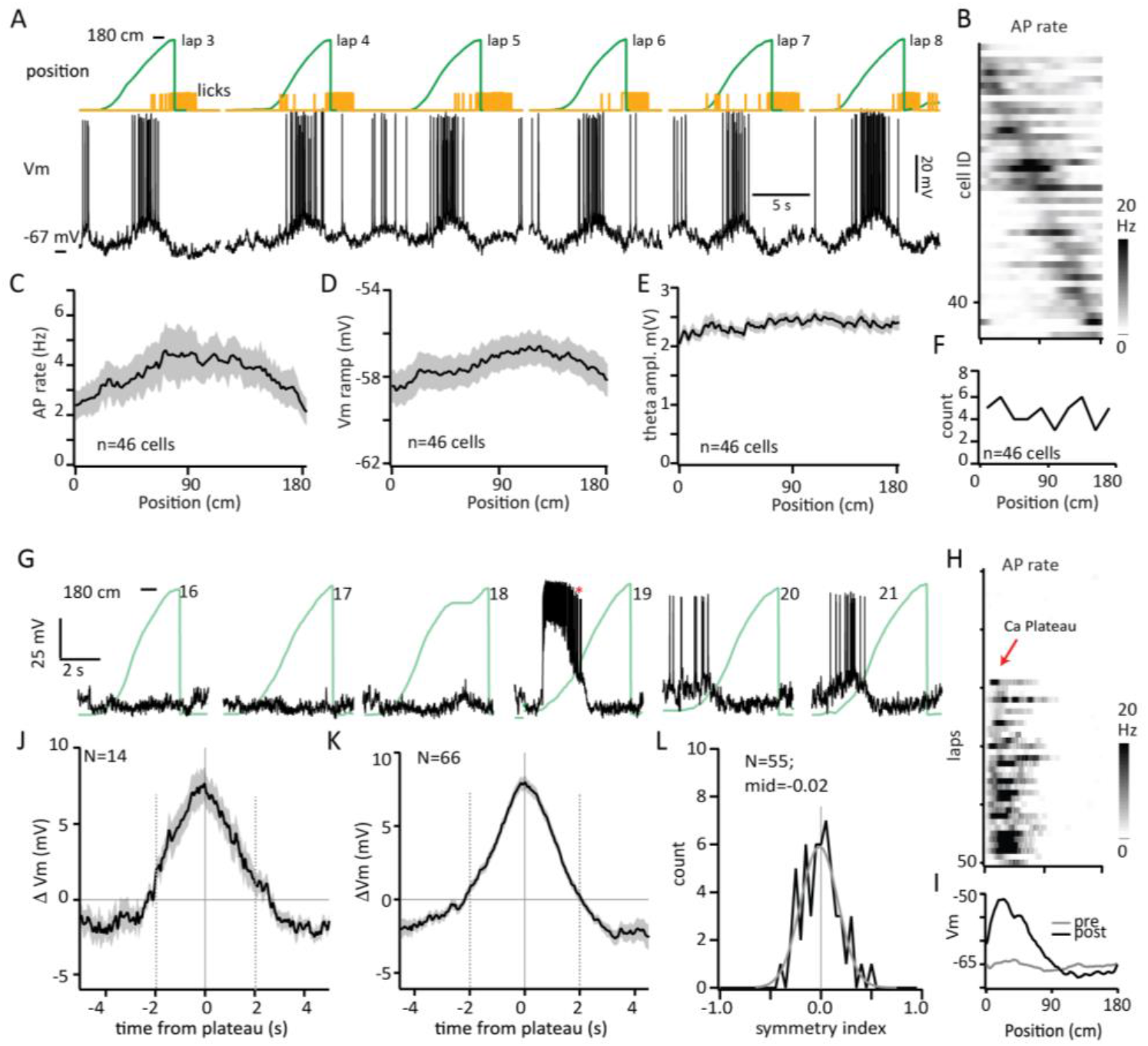
Symmetric bidirectional BTSP underlies CA3 PFs. (**A**) Membrane potential (Vm; black), mouse position on track (green) and mouse licking (yellow) for six laps. Vm during prolonged standing between laps are removed (breaks). (**B**) Heat map of PF firing for all PCs. Spatial profile of average (**C**) AP rate, (**D**) Vm ramp, (**E**) Vm theta during running for all PCs. (**F**) Histogram of PF peak location for all PCs. (**G**) Vm (black) and position (green) traces from CA3 neuron for multiple trials showing abrupt PF formation following naturally occurring plateau. (**H**) Heat map of AP firing rate and (**I**) average Vm ramps from laps before (pre) and after (post) plateau from PC in (G). The difference between Vm ramp after and before plateau (ΔVm) for the population with (**J**) naturally occurring plateau and (**K**) experimentally-induced plateau. (**L**) Histogram of symmetry index for population.

Next, we observed in a small fraction of principal cells (14/185) the initiation of long-duration Ca-plateau potentials or even trains of such plateaus in neurons lacking a PF or outside of an existing PF in PCs (ADP duration increased ∼6 fold over within-field ADPs; Fig. 1, G and H, and fig. S3). The initiation of long duration plateau immediately altered the Vm depolarization profile in these cells through bidirectional effects on Vm (Fig. 1, I and J, and fig. S3). Although this Vm plasticity was reminiscent of BTSP induced PF formation in hippocampal area CA1^25-33^, the resulting change in Vm (ΔVm) appeared to be symmetric in time for CA3 neurons (Fig. 1J and fig. S3). This is in contrast to BTSP in CA1 which has a markedly asymmetric time course.

To more thoroughly examine this uniquely symmetric form of PF plasticity in CA3 we injected large amplitude currents into a group of neurons at locations on the track over multiple trials (1 nA for 500 ms; n=78; average 5.8±0.1 induction trials; fig. S4, A to E). These “induced” plateaus again produced a bidirectional Vm plasticity in the majority of neurons (66 of 78) whose time-course symmetrically extended ∼10 s around the induced plateaus (Fig. 1, K and L, and fig. S4, F to K). Although the time course of plateau-induced PF plasticity was different in CA3, other properties were similar to BTSP in CA1 (fig. S5). Together these data indicate that a uniquely symmetric form of bidirectional BTSP underlies PFs in CA3. This symmetric form contrasts the highly asymmetric BTSP time-course observed in CA1. Symmetric synaptic plasticity rules are generally thought to promote the formation of stable attractors as they provide positive feedback that is directionally unbiased with respect to movement of the animal (fig. S1).

### Inputs involved in CA3 PF formation

CA3 PFs have been hypothesized to be formed through synaptic plasticity selectively at the various excitatory inputs to CA3 principal neurons including grid cell input from EC, mossy fiber input from the DG and among the recurrent CA3-CA3 connections as well^8-19,34-39^. Thus, we next questioned what synapses were being adjusted by BTSP to produce PFs in area CA3. To examine this we independently inhibited each of the excitatory input pathways to CA3 pyramidal neurons using optogenetic techniques (figs. S6 and S7). First, we inhibited CA3 itself by expressing ReaChR in PV+ interneurons and bilaterally directing excitation light to CA3 via chronically implanted optical fibers (fig. S6A). To test the role of CA3-CA3 recurrents we briefly inhibited CA3 activity just before the position of the BTSP inducing current injections (2.6±0.5 s, 67.1±1.8 cm; n=7) (Fig. 2A). This inhibition caused an approximately 7 mV hyperpolarization during the light (−7.5±1.2 mV) and dramatically impacted the shape of the resulting change in Vm compared with control animals (fiber implanted but lacking expression) that were also given a similarly sized Vm hyperpolarization via current injection (−7.8±2.0 mV; n=5) (Fig. 2, A to D, and fig. S6B). Inhibition of CA3 input significantly reduced the amount of BTSP induced potentiation at the location of the manipulation (i.e. before the plateau induction) and this caused a highly asymmetric shaped BTSP induced Vm change (Fig. 2, D and E, and fig. S8, A to C). This is consistent with the idea that, since inputs are required to be active for their weights to be altered, any manipulation that reduces the number of active inputs will result in less total synaptic potentiation (Fig. 2, F to I) and strongly indicates a role for CA3 recurrent inputs in PF formation^25,27,40^.

**Fig. 2.**
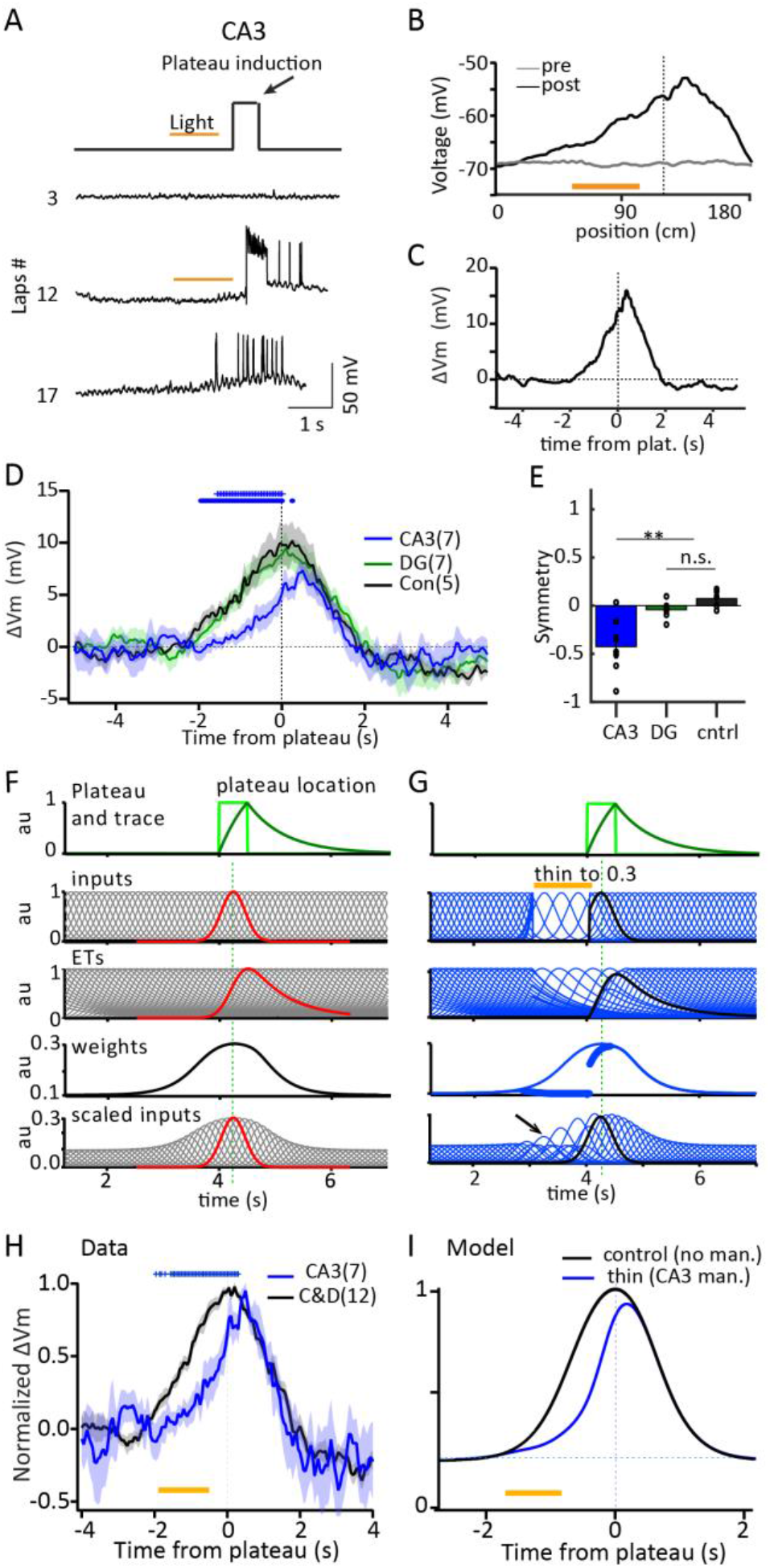
Involvement of CA3 and DG input pathways in PF formation. (**A**) Schematic (upper) and Vm traces for laps before, during and after induction with CA3 inhibition. (**B**) Vm ramps for laps before (gray) and after (black) plateau induction and (**C**) difference between Vm ramp after and before plateau (ΔVm) for neuron in (A). (**D**) Average ΔVm for populations showing impact of pathway inhibition on BTSP induction profile. All bins with significant difference are shown for two-tailed unpaired students t-test (p<0.05). Upper blue symbols, CA3 vs control; lower blue symbols, CA3 vs DG. (**E**) Average symmetry index for each group. Values from individual neurons shown. Symbols indicate significant difference (p<0.05 for two-tailed unpaired students t-test) for CA3 versus DG and CA3 versus control, and not significant for DG versus control. (**F**) Plateau voltage (light green) and hypothetical biochemical filter of plateau potential (plateau trace; dark green), control model signals (gray; top to bottom); input signals (inputs), hypothetical biochemical filter of synaptic input (eligibility traces; ETs), BTSP induced weight distribution (weights), inputs scaled by weights (scaled inputs). Red signals are those associated with the centermost input. (**G**) Same signals for “CA3 manipulation” model (blue) that simulated a 70% reduction in input numbers for one 1 s (thin to 0.3; yellow bar). Larger blue circles are weights affected by manipulation. Arrow indicates region where weights were affected by manipulation. Scaled inputs are control input sequence scaled by weights resulting from manipulation. Black traces are signals associated with the centermost input. (**H**) Average peak scaled ΔVm for populations showing the impact of pathway silencing on BTSP induction profile shape. Control and DG groups are combined (black). All time bins with significant difference are shown for two-tailed unpaired students t-test (p<0.05). (**I**) Simulated Vm ramps produced by summation of scaled inputs resulting from control (black) or CA3 manipulation (blue) conditions. Every 10^th^ signal shown in (F) and (G).

We next examined the impact of inhibiting DG input on the shape of the BTSP induced Vm change (DG expression of archaerhodopsin with bilateral optical fibers including one directed at CA3 recording site; figs. S6C and S7). In this case, we also included appropriately sized hyperpolarizing current injections (−10.4±1.2 mV; n=7) as this inhibition did not itself cause an observable Vm hyperpolarization. Inhibiting DG in this way had no impact on the profile of the resulting Vm change, which was basically symmetric around the induced plateaus (Fig. 2, D and E, and fig. S8, D and E).

Finally, we implanted optical fibers in the EC in mice expressing ReaChR in PV+ interneurons to test the role of excitatory input from this brain region (ipsilateral inhibition only; n=13; Fig. 3A and fig. S6E). Unexpectedly, we observed that the effect of this manipulation was basically the opposite of that produced by inhibiting CA3 input. That is, the Vm potentiation present when the EC was inhibited (i.e. before the plateaus) was enhanced and that after the plateau was decreased (Fig. 3, B to D). This produced an asymmetry in the opposite direction of that observed for CA3 inhibition (Fig. 3, D and E, and fig. S8F). This asymmetry can be explained if excitation from EC was responsible for linking the movement of the CA3 activity bump with the movement of the animal in the environment (Fig. 3, F to I). In this case, CA3 activity during the light activation would persist for longer than normal (fig. S9, B and D) and the increased amount of synaptic input delivered by this persistent activity would elevate the amplitude of the associated eligibility traces (ETs) (see arrow in Fig. 3G). These larger ETs would lead to more potentiation of the persistent inputs due to a greater amount of overlap with the plateau trace^25,27,40^ (PT; see Methods). In addition, the prolonged activity during the manipulation will accordingly delay the arrival of subsequently active inputs reducing their potentiation due to less overlap between the activity traces (see highlighted waveforms in Fig. 3, F and G). Altogether the total final effect is a leftward shift in the induced Vm change (ΔVm) (Fig. 3, H and I).

**Fig. 3.**
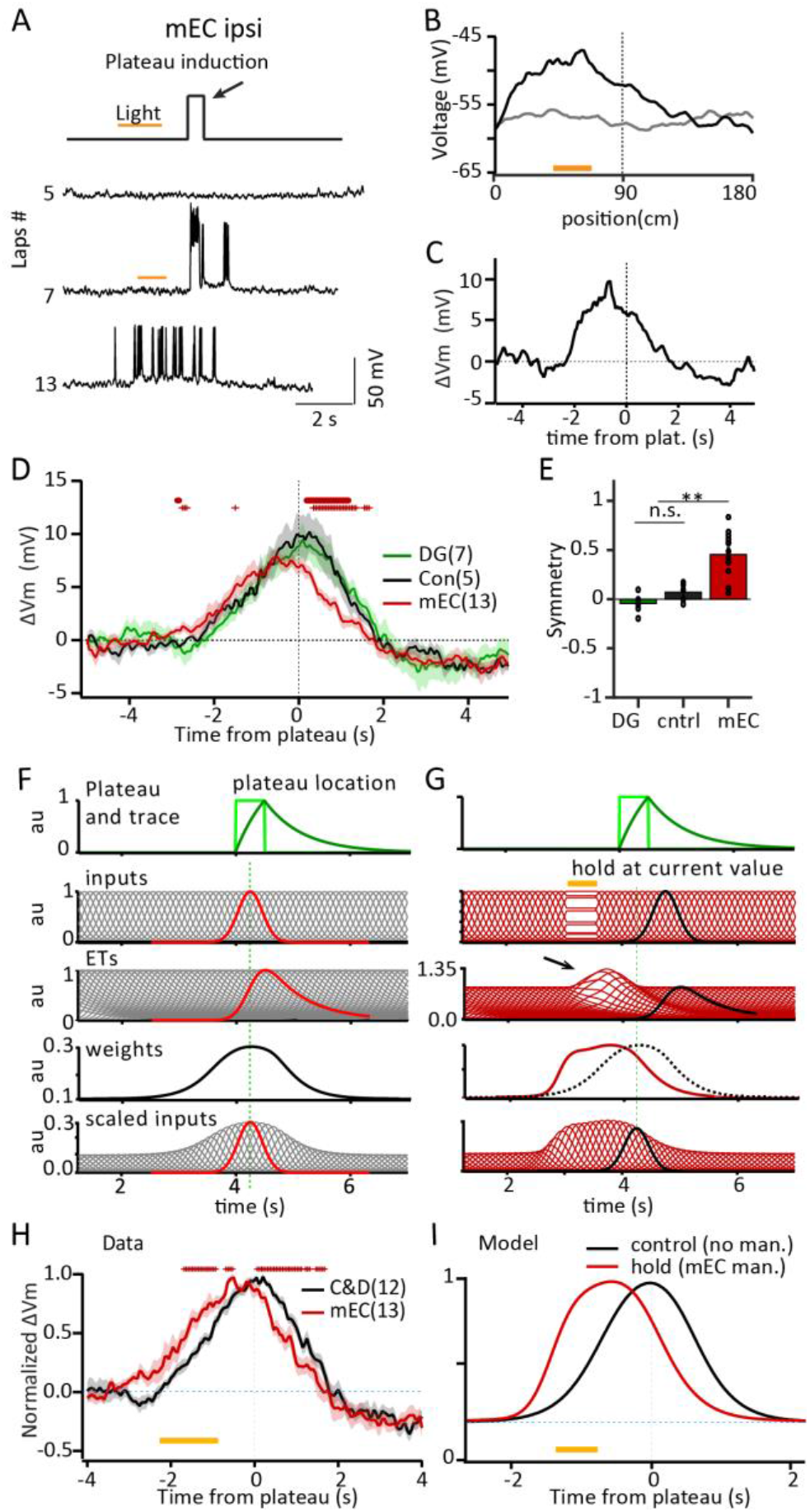
Involvement of EC input pathway in PF formation. (**A**) Schematic of optogenetic inhibition (upper) and Vm traces for laps before, during and after induction with EC inhibition. (**B**) Vm ramps for laps before (gray) and after (black) plateau induction and (**C**) difference between Vm ramp after and before plateau (ΔVm) for neuron in (A). (**D**) Average ΔVm for populations of neurons showing the impact of pathway silencing on BTSP induction profile. All time bins with significant difference are shown for two-tailed unpaired students t-test (p<0.05). Upper red symbols, EC vs control; lower red symbols, EC vs DG. (**E**) Average symmetry index for each group. Values from individual neurons shown. Symbols indicate significant difference (p<0.05 for two-tailed unpaired students t-test) for EC versus DG and EC versus control, and not significant for DG versus control. (**F**) Plateau voltage (light green) and plateau trace (dark green), control model signals (gray; top to bottom); input signals (inputs), eligibility traces (ETs), BTSP induced weight distribution (weights), inputs scaled by weights (scaled inputs). Red signals are those associated with the centermost input. (**G**) Same signals for “EC manipulation” model (red) where all inputs were held at their current amplitude for 0.5 second to simulate a hold of activity bump (yellow bar). Black signals are those associated with the centermost input. Dashed weights trace is from control. Scaled inputs are control input sequence scaled by weights resulting from manipulation. (**H**) Average peak scaled ΔVm for populations of neurons showing the impact of pathway silencing on BTSP induction profile shape. Control and DG groups are combined (black). All bins with significant difference are shown for two-tailed unpaired students t-test (p<0.05). (**I**) Simulated Vm ramps produced by summation of scaled inputs resulting from control (black) or EC manipulation (red) conditions. Every 10^th^ signal shown in (F) and (G).

In summary, only CA3 inhibition was associated with a decreased level of BTSP induced synaptic weight potentiation during the manipulation, while inhibition of the other pathways had either no effect or caused a shift in the BTSP induced Vm change. We interpret these data to indicate that it is BTSP at CA3-CA3 recurrent synapses that is responsible for the PF firing of CA3 PCs. Thus, new PF formation or modification of existing PFs during learning is produced by an adjustment of the synaptic weights of other currently active CA3 PCs.

### Inputs involved in CA3 PC activity updating

To further explore the above hypothesis that EC input is required for updating CA3 PF activity, we examined how inhibition of the different input pathways affected the Vm of CA3 neurons already expressing PFs (either natural or experimentally induced). Inhibition of CA3 had only a transient impact on Vm ramps such that the Vm hyperpolarized during the light but immediately returned to control values after light cessation (Fig. 4 and fig. S9, C and E). On average DG inhibition had no impact on the Vm ramp either before or after the light activation (Fig. 4 and fig. S9E). In contrast to the above effects, EC inhibition caused the Vm to essentially persist at its current level during the light, in effect causing a net hyperpolarization with respect to control laps (n=13 cells with ipsilateral and 3 cells with bilateral inhibition; Fig 4 and fig. S9, D and E). Following termination of the light the Vm slowly recovered its normal trajectory but with a delay that produced a shift in the Vm ramp after the light activation. The magnitude of this shift was correlated with the size of the light effect (Vm hyperpolarization; Fig. 4, E to G) but uncorrelated with any modest changes in animal running (Fig. 4H). These EC manipulation results provide further evidence that direct EC input to CA3 is necessary for a bump of activity in area CA3 to advance appropriately with the movement of the animal. Altogether the above data suggest that a symmetric form of BTSP at CA3-CA3 synapses forms neuronal attractors, the activity of which are updated according to the animal’s behavior by an external input from the EC.

**Fig. 4.**
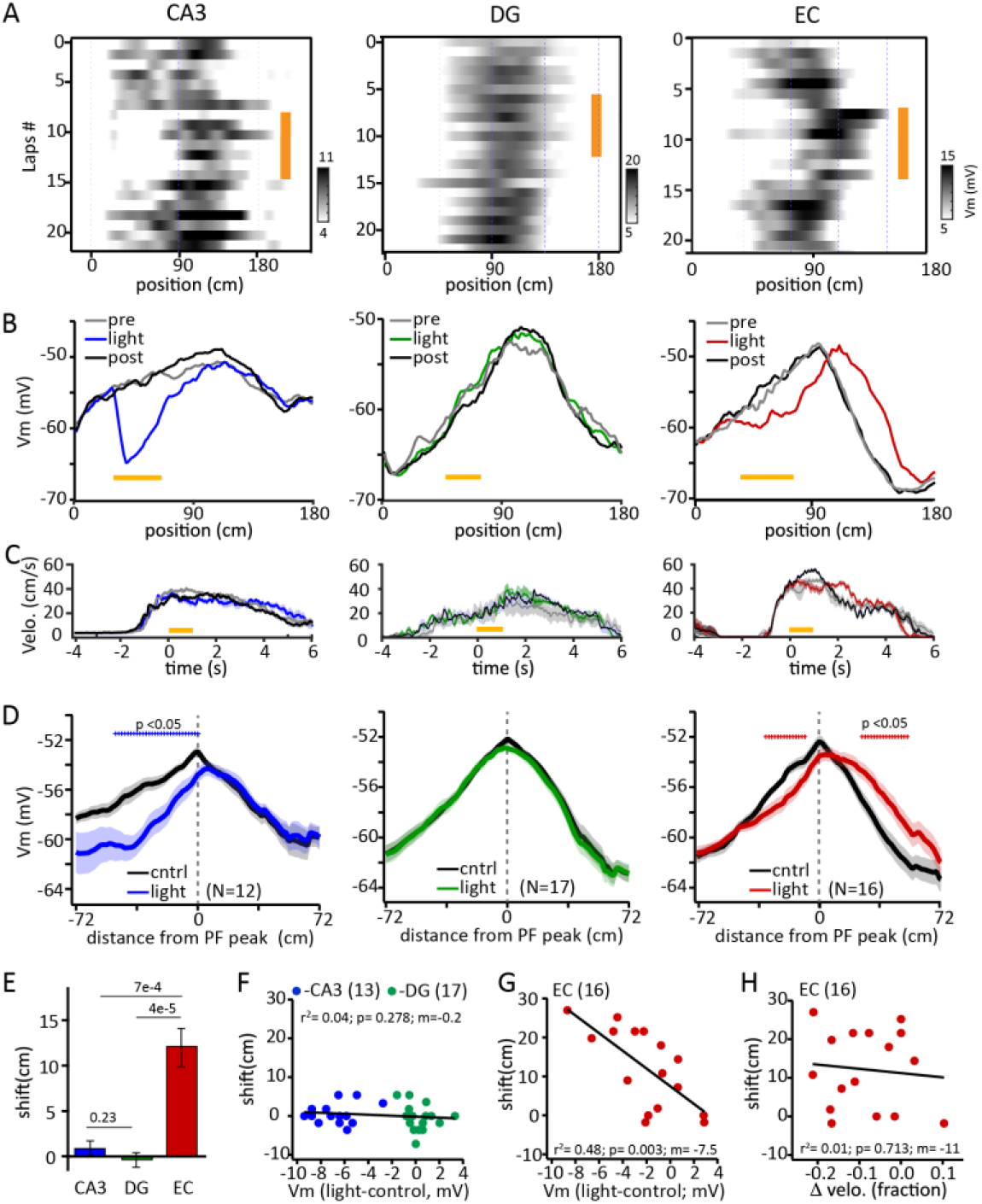
Involvement of different input pathways in PF activity. (**A**) Spatial profile of Vm ramps for listed trials. Yellow bar indicates trials where light was active to inhibit either CA3, DG or EC, from left to right. Vm ramps are baseline corrected by subtracting the mean value of 1st 5 bins. (**B**) Vm ramps for laps before, during and after light activation for different groups. Note the pause and subsequent shift in Vm ramp for EC inhibition. (**C**) Average velocity traces for the same trails shown above. (**D**) Group averages of Vm ramp for control trials (average of pre and post light trails, black) and manipulation trials (CA3, blue; DG, green; EC, red). Vm ramps are aligned to peak position on control trials. (**E**) Average level of Vm ramp shift after light activation (values for individual neurons shown in F and G). *p* values for two-tailed unpaired students t-test are shown. (**F**) Relationship between Vm ramp shift and impact of inhibition on Vm for CA3 and DG groups. **(G)** Relationship between Vm ramp shift and impact of inhibition on Vm for EC group. (**H**) Relationship between Vm ramp shift and impact of inhibition on running velocity.

### Attractor network model and theoretical capabilities

We next tested this hypothesis in a recurrent network model where different subsets of neurons emitted plateau potentials uniformly across all locations of an environment during simulated navigation. A symmetric BTSP plasticity rule was used to potentiate synapses between neurons emitting plateaus close in time and de-potentiate synapses between neurons whose plateaus were farther apart (fig. S10D). This rule produced a symmetric synaptic weight profile with spatially localized excitation (Fig. 5A). In a network that included uniform inhibitory feedback the bump of activity resulting from the network dynamics could be controlled by an external input. Indeed, once the external input was removed from the network, the attractor dynamics kept an activity bump stable at the last visited location, consistent with the above experimental observations (Fig. 5B). Thus, the unique symmetric time course of BTSP observed in CA3 was able to produce continuous attractor dynamics that were effectively controlled by an external input similar to that provided by the EC.

**Fig. 5.**
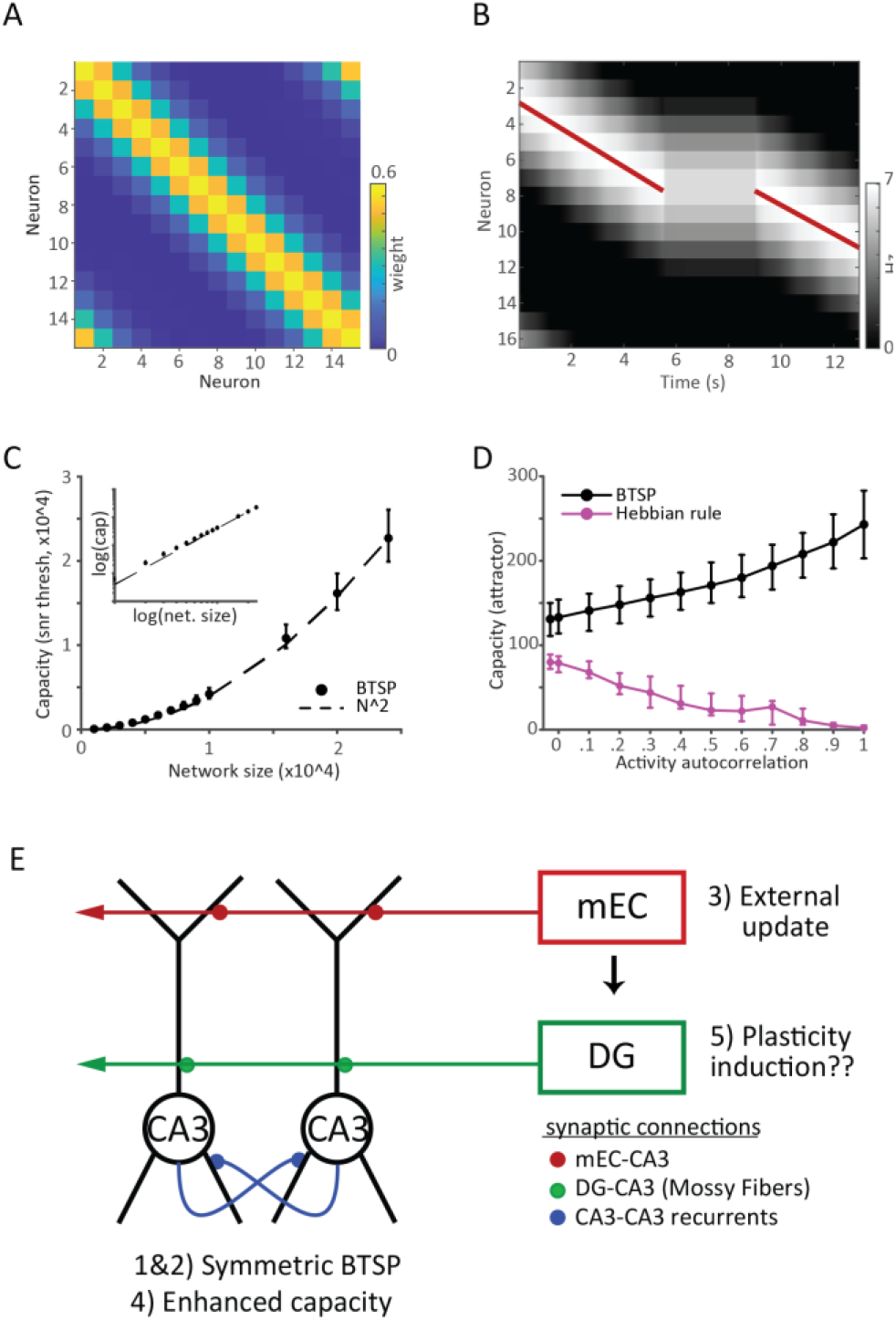
Attractor dynamics in recurrent neural networks with BTSP. (**A**) Synaptic weights learned between plateau-emitting CA3 model units after ten simulated laps on the belt, in the presence of ongoing BTSP. (**B**) A spatially localized input (red line) drives a bump of activity in the network. Upon input removal, the bump persists at the last visited location. (**C**) Optimal storage capacity scaling. Capacity is estimated from signal-to-noise ratio in networks of increasing size N, with sparse random uncorrelated pre- and postsynaptic activity and plateaus. The probability of having activity or plateaus in any given neuron at any given time is proportional to log(N)/N. The network’s memory span grows approximately quadratically with system size (N^2^/log(N)^2^, logarithmic scale plot in the inset). (**D**) BTSP is robust against temporal correlations in activity, outperforming rules relying solely on simultaneous pre- and postsynaptic activity. Values plotted are capacity where probability of correct decoding drops to 0.75 (lower bar), 0.50 (data point), 0.25 (upper bar). Parameters for (A-D) in Methods. (**E**) CA3 circuit schematic showing various excitatory synaptic input pathways and their approximate locations on a pair of CA3 pyramidal neurons compromising a single attractor. Numbered text is hypothesized roles for each input according to questions posed in introduction

Neuronal networks possessing attractor dynamics such as those observed in the above experiments and model are effective at storing memories and are found in many theories of hippocampal mnemonic function^8-14^. The actual storage and retrieval capabilities of such attractor networks are shaped by the implemented synaptic learning rules^41-48^. Therefore, we assessed the memory capacity of a network endowed with a symmetric form of BTSP by measuring its ability to retain previously stored input patterns as additional patterns were presented (fig. S10, A to C). The measured signal-to-noise ratio (SNR; see Methods and Supplementary Materials) exhibited an exponential decay with increased numbers of patterns and memory capacity was determined as the number of patterns producing an SNR larger than a certain threshold (fig. S10B; see Methods and Supplementary Materials)^43^. As we increased the size of the network, we observed an approximate quadratic scaling of memory capacity, which is optimal from an information-theoretic standpoint (Fig. 5C; see Methods and Supplementary Materials)^41,43^. Furthermore, we observed a surprising increase in the number of attractors that can be stored in the network as temporal correlations in the activity patterns were increased (Fig. 5D). This robustness to temporal correlations is in stark contrast to that observed in a Hebbian type rule (Fig. 5D, same-time rule curve; fig. S10C) and is mediated by the more targeted depotentiation present in BTSP. Here depotentiation is more targeted as it acts only to reduce interference between a specific activity pattern and another highly correlated pattern that occurred in the recent past (figs. S10H and S11). These results demonstrate the efficacy of BTSP in generating large numbers of attractor states within recurrent neural networks, and the advantage of BTSP compared to traditional Hebbian plasticity.

## Summary and conclusions

Together the above data, theory, and simulations support the idea that hippocampal area CA3 exhibits attractor dynamics and point to cellular and circuit mechanisms that effectively mediate this activity (Fig. 5E). First, the density of CA3 PFs is uniform across the environment providing the flat energy landscape needed for stable dynamics. Second, the synaptic plasticity responsible for PF activity is a version of BTSP that has a symmetric time-course. In addition, this plasticity adjusts the synaptic weights among the CA3-CA3 recurrent pathway allowing these synapses to provide the positive feedback required for stable persistent activity among like-tuned PCs. Finally, in the attractor framework this stable activity bump requires an external input to link the movement of neuronal activity to that of the animal^9,10,13,14,16^ and this association appears to be mediated by the direct excitatory input from the EC. Thus, path integration related activity in the EC may be involved in the selection of which attractors will become active through what is subsequently an essentially self-organizing process. The role of EC-CA3 synaptic weight adjustments in this process remains an open question.

In the above scheme, learning of a new environment and the stable storage of relevant learned information involves a reshaping of current CA3 PC activity, as opposed to, for instance, any transformation from grid cells to place cells^34,46-51^. Thus, new CA3 PFs will be formed using the tuned activity of other CA3 PFs or plateaus via BTSP modification of synaptic weights between them. We provide theoretical analyses that indicates BTSP can effectively mediate such online learning in a recurrent network. Indeed, we found that learning and memory formation using this type of plasticity outperforms more standard correlation-based forms of plasticity. The effectiveness of our BTSP model in recurrent networks is partly based on the presence of sparse plateaus in the network^28,32,52^, which in CA3 are comparable in duration with PF activity (fig. S3). This sparsity, akin to the sparse coding hypothesis for neural activity^41,43,53,54^, helps to efficiently represent and store information with minimal overlap between representations. Moreover, the depotentiation component of BTSP will transform similar input patterns into more distinct output patterns, which will contribute to pattern separation by preventing interference between similar patterns at the level of synaptic dynamics. Together the sparsity and targeted depotentiation present in BTSP enhance the ability of hippocampal area CA3 to create unique representations and therefore increase memory storage and retrieval capacity. This ability could also make BTSP a key component in global remapping, a process in which the hippocampus rapidly reorganizes its activity patterns in response to changes in the environment or task^55,56^.

The evidence presented here indicates that BTSP in the CA3 recurrent network near optimally forms unique neuronal representations with features associated with attractor dynamics, including the ability to link activity updating with the behavior of the animal via an external input from the EC. Similar plasticity rules and circuit mechanisms could also be highly effective in other recurrent networks, such as those found throughout the neocortex. Overall, the above observations advance our understanding of the mnemonic function of the hippocampus by providing evidence that unique instantiations of BTSP and activity updating circuit mechanisms within hippocampal area CA3, a brain region long associated with episodic memory, efficiently and effectively support a form of network dynamics that remains the most compelling framework for memory storage and retrieval in brains.

## Acknowledgments

We thank R Chitwood for technical support, A. Roxin, G. Buzsaki, H. Inagaki, N. Brunel, and N. Li, and for useful comments on the manuscript. This work was funded by HHMI (JCM, SR), the Cullen Foundation (JCM) and HHMI via Life Sciences Research Foundation (YL).

## Author contributions

Conceptualization: YL, JJB, SR, JCM

Physiological recordings: YL

Physiological data analysis: YL, JCM

Computational modeling and Theory: JJB, SR

Writing, review & editing: YL, JJB, SR, JCM

## Data and materials availability

The data supporting this study’s findings are available from the corresponding author upon request. The code that supports the results of this study will be made available via a GitHub repository.

