## supplementary materials for "Mechanisms of memory storage and retrieval in hippocampal area CA3"

**The PDF file includes:**

Materials and Methods
Supplementary Text for BTSP model
Figs. S1 to S11
References (57-64)

#### Materials and Methods

##### Animals and procedures

All experimental methods were approved by the Baylor College of Medicine Institutional Animal Care and Use Committees (Protocol 15-126). All experimental procedures in this study, including animal surgeries, behavioral training, treadmill and rig configuration, and intracellular recordings, were performed identically to a previous detailed report[25-27] with the exception that area CA3 was targeted instead of CA1.

Wild type C57BL6 (*WT*) male mice aged 10-14 weeks obtained from the Baylor College of Medicine Center for Comparative Medicine or Jackson Laboratory were used for all non-manipulation experiments. Different transgenic mice in 10-18 week-old of either sex were used for different sets of optogenetic experiments: *PV-cre::ReaChR* was used for CA3 and EC silencing, non-crossed *ReaChR* was used as the control group, *Rbp4-cre::Ai35* and *Rbp4-cre::ReaChR* were used for DG silencing and activation, respectively.

Craniotomies above the dorsal hippocampus for simultaneous whole-cell recordings and local field potential (LFP) recordings, as well as affixation of head bar and optical fiber implants were performed under deep anesthesia. The coordinates for hippocampal CA3 recording were (in mm): AP: 1.85, ML: 2.35 (fig. S2, A and B). The optical fibers (200 um in diameter, 0.5 N.A., Thorlabs) were implanted bilaterally at 0.15 mm (AP), 3.0 mm (ML), 40° to the horizontal, 35° to the sagittal, and the depth is ~1.9 mm along the axis of the fiber for CA3 and DG manipulations as well as control groups (fig. S6, A to D). For the EC manipulations, the fibers were implanted ipsi-laterally/bilaterally at 4.8 mm (AP), 3.4 mm (ML), 10° to the coronal and the depth is 1.1 mm along the axis of the fiber (fig. S6E).

Following a week of recovery, animals were prepared for behavioral training with water restriction, handling by the experimenter, and addition of running wheels to their home cages. Once the training started, water was given only during the experiment and in the end of the day, and specific care was taken to keep the animal body weight above 80 % of the weight before water restriction. Mice were trained to run on the cue-enriched linear treadmill for a 10 % sucrose reward delivered through a licking port once (~3.3 ul) per lap (~187 cm). All mice were reared on a reversed 12/12 hr light/dark cycle.

*In vivo* intracellular electrophysiology

To establish whole-cell recordings from CA3 pyramidal neurons, an extracellular LFP electrode was lowered into the dorsal hippocampus using a micromanipulator until prominent theta-modulated spiking and increased ripple amplitude was detected again after transiting the CA1 pyramidal layer, usually to a depth of 1.8-2.0 mm. Then a glass intracellular recording pipette was lowered to the same depth while applying positive pressure. The intracellular solution contained (in mM): 134 K-Gluconate, 6 KCl, 10 HEPES, 4 NaCl, 0.3 MgGTP, 4 MgATP, 14 Tris-phosphocreatine, and 0.2 % biocytin. Current-clamp recordings of intracellular membrane potential ($V_m$) were amplified and digitized at 20 kHz, without correction for liquid junction potential.

A MATLAB GUI interfaced with a custom microprocessor-controlled system for position-dependent reward delivery and intracellular current injection. Animal run velocity was measured by an encoder attached to one of the wheel axles.

For plateau induction, a position-dependent step current (1 nA, 500 ms, usually 5 laps) was injected into the recorded cells. In a subset of cells where resting Vm was greater than ~-65 mV an additional small DC current (<50 pA) was also given to increase the probability of inducing plateaus. A 100 Hz current train (1 nA, 500 ms, 5 laps) instead of the step current was injected to induce APs in some experiments (fig. S5, F to J).

Place field analysis

To analyze the spatial location of AP rate, the spatially binned AP rate was determined (AP#/time in bin) using 100 equally sized spatial bins (~1.80 cm) for each trial and averaged over the duration of the recording. To analyze $V_m$ ramps, raw $V_m$ traces were median filtered to remove APs, baseline corrected by subtracting the difference in the recorded AP threshold and -50 mV and spatially binned and averaged as above. To analyze theta frequency Vm oscillations, the Vm ramp were first bandpass filtered (2-10 Hz) and then the amplitude of the theta-band oscillations was determined from a Hilbert Transform of these traces. In some cases (Fig. 1), the average spatially-binned AP rates, Vm ramps and theta amplitude traces were further smoothed using a boxcar of 5 bins. The spatially-binned AP rates in the heat map shown in Fig. 1E were smoothed with a gaussian (36 cm). Spontaneous naturally-occurring plateau duration was estimated as the full width at plateau detection threshold Vm value (-35 mV).

To determine the plateau-induced Vm change 5-10 trials before the plateau trials were averaged as a “before” trace. 5-10 trials after the plateau were also averaged as an “after” trace and this was subtracted from the before trace to produce a difference (ΔVm). For naturally occurring plateau the time base (in 100 spatial bins) was produced by determining the time required for the animal to run from a given track location to the location of the ΔVm peak on the trial where the natural plateaus occurred. For induced plateau the time base was determined as the minimum run time from the middle of the plateau-inducing current injection to a particular location (in 100 spatial bins) as calculated from all induction laps. For this a single composite position versus time trace that represented the shortest induction lap in time was constructed by taking the minimum time delay to plateau midpoint for each spatial position across all induction laps (fig. S4C).

$V_m$ ramp half-width (fig. S5, D, I and M) was calculated from the $\Delta V_m$ traces as the time (s) or distance (cm) between the plateau and the final return of $\Delta V_m$ to 15 % of max). This value was halved if both sides of the ΔVm trace returned to the minimum value, if only one side returned (because the end of the track was reached before) then this single value was used. The average velocity was calculated from the composite induction lap as the mean velocity of the mouse for the distance covered by the half-width measure (fig. S4). The symmetry of the ΔVm trace in time was determined from the relative difference in the positive areas of the two sides of the curve, (positive time side - negative time side) / total.

Optogenetic manipulations

Optical fibers were coupled to an external fiber using standard FC connectors via the mental sleeve and then connected to two 595 nm LEDs (Thorlabs) or a 594 nm laser (Cobolt) (only used in part of DG silencing experiment (fig. S7, C to H)). The maximum power (~2.5 mW for LED and ~23 mW for laser) was used in all the optogenetic experiments except the ipsi-EC manipulation. For the ipsi-EC manipulation, the power (0.8-2.5 mW) was adjusted accordingly so as to not affect the animal’s behavior. A 40 Hz sine wave light stimuli, generated by an arbitrary waveform generator (Tektronix), was given to activate the PV positive interneurons expressing ReaChR, and a constant step light stimuli was used to silent/activate the granule cells expressing Arch/ReaChR. The light delivery (position-dependent onset, distance-/time-based duration) was controlled by the custom microprocessor-controlled system.

Comparison of optogenetic manipulations with other studies is difficult due to different recording conditions, manipulation durations and behavior[58-64]. Most studies have recorded in CA1 (but see 58,59) while inhibiting EC or CA3. The most common effect of inhibiting EC is an induction of global remapping of PF activity. Since this is a population phenomenon and we are recording from single cells it is difficult to compare our results (ie a shift in PF activity) with these other studies. However, as far as we can surmise no other studies have reported observing such a shift in PF activity.

The level of Vm ramp shift produced by optogenetic manipulations was quantified as the amount of shift required to minimize the difference between the decay portion of the Vm ramp (from peak to end of track) for test and control traces. The delta velocity was determined as the difference in the mean velocities for the time period during the light activation for test and control laps (velocity traces referenced to the beginning of light activation; Fig. 4C) and quantified as a fractional change (ΔVel/average Vel). Net effect of light on Vm was quantified as the difference in the average Vm during the light and Vm for the same spatial bins of control traces from spatially-binned Vm ramps. Negative values represent net Vm hyperpolarization (i.e. less depolarization) during the light.

Histology

After the end of recordings, a LFP pipette loaded with DiI (saturated in DMSO) was lowered to 200 um above the CA3 pyramidal layer and a positive pressure (2-4 psi) was given to fill the pipette track with DiI. Mice were perfused transcardially with PBS and then with 4 % PFA in PBS after deep anesthesia. The brain was dissected out and further post-fixed in 4 % PFA overnight, and then stored in PBS at 4 °C. The fixed brain was cut into 100 um thick coronal/sagittal sections with a vibratome. Sections around the recording site/optical fiber (with DiI/fiber track) were harvested and then incubated with fluorophore-conjugated streptavidin (1:1000, Invitrogen) against biocytin for 2 h at room temperature. The fluorescence images were acquired using Leica SP8X confocal microscopy (fig. S2, A and B) or Zeiss AXIO Zoom.V16 stereo microscopy (fig. S6).

Quantification and statistical analysis

Statistical details of experiments can be found in the figure legends. Unless otherwise specified, measured values and ranges reflect mean ± SEM. Significance was defined as $p<0.05$. Sample sizes were not determined by statistical methods, but efforts were made to collect as many samples as was technically feasible. No data or subjects were excluded from any analysis.

#### Computational modeling

##### *Schematic BTSP model for Figs. 2 and 3*

CA3 inputs are represented by a set of 1000 gaussian functions (ampl=1; SD=0.5s) whose midpoints were 10 ms apart. These functions are meant to mimic a sequence of tuned inputs moving in time from 0 to 10 s. Eligibility traces (ET) are convolutions of each of the gaussian inputs with an exponential function ($\tau$=0.67 s). These signals simulate a biochemical filter of the input. Plateau traces (PT) are the convolution of the plateau voltage signal (300 ms step) with an exponential function ($\tau$ =0.67 s). This signal simulates a biochemical filter of the plateau. Symmetric BTSP is produced by using activity filters (ET and PT) with the same time constant. Each input can have a weight from 0 to 1 and are initialized to 0.1. To simulate learning via BTSP, weights are changed according to a BTSP learning rule that only contained a potentiation component $\Delta W= (1-w_0)\, q_+(ET \cdot PT)$, where $w_0$ is the initialized weight, $q_+$ was a sigmoid function (slope=0.1; midpoint=0.25). Inputs are then scaled by the weights and summed to simulate a Vm ramp.

To mimic the optogenetic inhibition of CA3 input during BTSP induction, two out of every three inputs were set to 0 either for the remaining duration of simulation if the inputs peaked during the manipulation or for the duration of the manipulation if they peaked after the manipulation (SFig 9c). The manipulation occurred just before the start of the plateau (3s to 4 s; plateau initiation at 4.25s) during the simulation run used to calculate $\Delta W$. The manipulated inputs generated ETs with reduced amplitudes and thus reduced weight changes. Standard input trains (ie without manipulation) were then scaled by the resulting weight distribution and summed (see fig. S9). To mimic the inhibition of EC input, inputs were held that their current value for 0.5 s just before the plateau and then allowed to continue by altering the gaussian function accordingly. These prolonged inputs produced elevate ETs that were used to calculate $\Delta W$. The resulting weight distribution was then applied to normal input sequences and these inputs were summed. To simulate the effect of such input manipulations on already formed "PFs" the same manipulations we applied to the inputs and a standard weight distribution was used to scale each of the inputs. These inputs were then summed.

*Binary model (Fig 5 C and D)*

To gain a deeper understanding of the storage and retrieval capabilities of a network endowed with BTSP under general conditions, we sought to simplify the underlying rule governing synaptic plasticity while preserving the essential aspects of BTSP. We report a complete derivation and mathematical analysis of this rule in Supplementary Information, but we briefly describe the main concepts and features of the rule below.

We discretized time into broad windows (~4s) and introduced binary signals for the neurons, which differed depending on whether the neuron was pre- or post-synaptic for a given synapse (fig. S10, A and B). The two most prominent synaptic changes in BTSP occur in the form of (1) potentiation in the presence of presynaptic activity/plateau and postsynaptic plateau occurring within ~2s of each other, and (2) depotentiation with temporally offset (from ~2s to ~4s) postsynaptic plateau and presynaptic activity/plateau. Further, by assuming that synapses are binary (being either in a potentiated or depotentiated state), we approximated BTSP as:

$$W(t+1) = W(t) + \big(1 - W(t)\big) q_{pre}(t) q_{post}(t) - W(t)\Big(q_{pre}(t) q_{post}(t-1) \; OR \; q_{pre}(t-1) q_{post}(t)\Big) \quad (1)$$

where $\mathrm{W} = 0{,}1$ represents whether the synapse is depotentiated or potentiated, respectively, $\mathrm{q_{pre}(t)} = 0{,}1$ represents whether activity or plateau occurred in the presynaptic neuron at time t and $\mathrm{q_{post}(t)} = 0{,}1$ represents whether or not a plateau occurred in the postsynaptic neuron at time t.

We next sought to eliminate the dependency on $(t-1)$ from the dynamics in Eq. 1, in order to allow an analysis of these dynamics in terms of a Markov chain. By introducing the variables $\alpha(t) = W(t) q_{pre}(t-1)$, $\beta(t) = W(t) q_{post}(t-1)$, the dynamics in Eq. 1 becomes:

$$
\begin{cases}
W(t+1) = W(t) + \big(1 - W(t)\big) q_{pre} q_{post} - W(t)\big(\beta(t) q_{pre} + \alpha(t) q_{post} - \alpha(t)\beta(t) q_{pre} q_{post}\big) \\
\alpha(t+1) = W(t) q_{pre} + \big(1 - W(t)\big) q_{pre} q_{post} - W(t)\big(\beta(t) q_{pre} + \alpha(t) q_{pre} q_{post} - \alpha(t)\beta(t) q_{pre} q_{post}\big) \\
\beta(t+1) = W(t) q_{post} + \big(1 - W(t)\big) q_{pre} q_{post} - W(t)\big(\beta(t) q_{pre} q_{post} + \alpha(t) q_{post} - \alpha(t)\beta(t) q_{pre} q_{post}\big)
\end{cases} \quad (2)
$$

This system exhibits an exponential memory decay (see SI). When presynaptic and postsynaptic activity and plateaus are temporally uncorrelated, the memory span of the system (the time constant of the exponential decay) is given by

$$
\tau_{uncorr} \approx \frac{1}{6 f_{pre} f_{post}} \quad (3)
$$

up to logarithmic corrections, where $f_{pre}$ is the probability of having activity or plateau presynaptically at each time step, and $f_{post}$ is the probability of having postsynaptic plateau. When presynaptic activity is correlated in time, we find that the memory span becomes:

$$
\tau_{corr} \approx \frac{1}{6 f_p^2 + 2(3 - c - c^2) f_a f_p} \quad (4)
$$

with $f_p$ representing the plateau probability, $f_a$ representing the activity probability, and $c$ represents the correlation of activity at adjacent timepoints. The capacity estimate based on the

SNR from network simulations reported in Fig. 5c had parameters $f_a = f_p = 10 \log N / N$ and $c = 0$.

*Network dynamics for storage capacity*

In Fig. 5D we built an attractor network using the weights learned from a recurrently connected network with global inhibition and thresholded activity described by the dynamics:

$$A_i(t+1) = \Theta\left(\sum_j (W_{ij} - \bar{W}) A_j(t) - \theta\right) \tag{5}$$

where $\vec{A}$ is the population activity, $W_{ij}$is the learned matrix, $\bar{W}$ is its average, $\Theta$ is the Heaviside function (1 when its argument is greater than zero and zero when the argument is less than zero), and $\theta$ is a threshold. The threshold was chosen to be $m + 3.5s$ where $m$ and $s$ are the mean and standard deviation of the distribution of inputs to units which are inactive $\sum_j W_{ij} q_{pre}^j \,| q_{post}^i = 0$. The attractor capacity was defined as the number of steady state activity patterns that the network can retrieve. To define successful retrieval, we built a decoder that chooses the pattern with maximal correlation to the steady-state attractor activity and compared this decoder's output to the pattern that seeded the attractor activity. The attractor capacity is then the number of patterns correctly decoded before the probability of correct decoding drops below 0.5. The results of varying $c$ with $N = 3000$, $f_a = f_p = 10 \log N / N$ are plotted in Fig. 5D.

*Network dynamics for continuous attractors*

For the results reported in (Fig. 5, A and B, and fig. S10D), we used the continuous version of the discrete BTSP model. To derive the model, we first compute the difference of each variable between time $t + 1$ and $t$ from Eq. 2. We then take the average across realizations of pre and postsynaptic signals In order to compute the average of the product $\alpha\beta$, we have to introduce an additional variable $\gamma = \alpha\beta$ and derive its dynamics. We finally approximate the finite difference with a time derivative. This results in the dynamics:

$$
\begin{cases}
\dot{W} = (1-W)q_{pre}q_{post} - \left(\beta q_{pre} + \alpha q_{post} - \gamma q_{pre}q_{post}\right) \\
\dot{\alpha} = -\alpha + W q_{pre} + (1-W)q_{pre}q_{post} - W\left(\beta q_{pre} + \alpha q_{pre}q_{post} - \gamma q_{pre}q_{post}\right) \\
\dot{\beta} = -\beta + W q_{post} + (1-W)q_{pre}q_{post} - W\left(\beta q_{pre}q_{post} + \alpha q_{post} - \gamma q_{pre}q_{post}\right) \\
\dot{\gamma} = -\gamma - q_{pre}q_{post}(\alpha + \beta - \gamma - 1)
\end{cases} \tag{6}
$$

In (fig. S10C) we estimated the change in synaptic weight by simulating these synaptic dynamics in the presence of Gaussian pre and postsynaptic signals, mimicking a postsynaptic plateau centered around $t = 0$, and presynaptic plateaus at $50$ equally spaced timing from $-4\ s$ to $4\ s$. We repeated this procedure for $50$ different initial values of the initial weight.

The Gaussian profiles had a standard deviation $\sigma = 0.75\ s$. For (Fig 5A) we modeled one plateau for each unit $i$ as baseline subtracted firing rate profiles described by Von Mises functions:

$$
\frac{e^{\kappa \cos(vt-\theta_i)} - e^{-\kappa}}{2\pi I_0(\kappa)} \tag{7}
$$

where $v = \pi$ is the velocity of the simulated animal in a circular environment (one lap in $2\ s$), $\kappa = 4$ is the concentration parameter, and $I_0(\kappa)$ is the zero order modified Bessel function of the first kind. Each of the $N = 16$ units in the network emitted a plateau around location $\theta_i$, which was uniformly spaced along the circular environment. The synaptic dynamics was simulated for ten laps, starting from a uniform initial condition for the weights, $w = 0.25$. We then used the resulting weight matrix $W_{ij}$ in a rate model dynamics

$$
\tau \dot{h}_i = -h_i + \frac{J}{N}\sum_j (W_{ij} - \bar{W})\phi(h_j) + I_0 + I_1 \cos\left(\theta_i - x(t)\right) \tag{8}
$$

where $\tau = 10\ ms$ is the integration time constant of the individual units, $J = 10$ is a scaling factor for the weights, $\bar{W}$ is the average of the weight matrix across all elements, $\phi()$ is the threshold-linear function, $I_0 = 2$ is a constant uniform input, and the last term described a tuned external input around the location of the animal $x(t)$ (red line in Fig. 5B). $I_1 = .25$ was the amplitude of the tuned external input. $I_1$ was set to $0$ to probe the existence of attractors in the network (Fig. 5B, time interval without the red line).

### Methods for BTSP models

#### Markov chains preliminaries

In the following we will consider plasticity rules whose dynamics can be described by a system of discrete-time equations for $d$ discrete variables $v_i(t), i = 1..d$

$$v_i(t+1) = F_i(v_1(t), .., v_d(t), q_1, .., q_s \tag{1}$$

where $q_i, i = 1..s$ are $s$ independent and uncorrelated random variables sampled at each time step from prescribed distributions, and the discrete functions $F_i(), i = 1..d$ determine the dynamics of the system.

We are interested in the Markov chain description of these dynamics. To do this we define the set of $n$ possible states that the dynamic variables $\vec{v}(t)$ can occupy as $\vec{V}^{(i)}, i = 1..n$. Note that the number of available states $n$ can be smaller than the number of possible words that can be obtained with $d$ discrete variables, due to the specific constraints introduced by the dynamics $F_i()$ (see examples in the following sections). The transition probability $M_{i,j}$ from state $j$ to $i$ can then be computed as

$$M_{i,j} = \sum_{\vec{Q}} P(\vec{Q}) \delta_{V_1^{(i)}, F_1(V_1^{(j)}(t),..,V_d^{(j)}(t), Q_1,..,Q_s)} \cdots \delta_{V_d^{(i)}, F_d(V_1^{(j)}(t),..,V_d^{(j)}(t), Q_1,..,Q_s)} \tag{2}$$

where $\delta$ is the Kronecker delta function and the sum is over all the realizations $\vec{Q}$ of the random variables $\vec{q}$, and $P(\vec{Q})$ is the corresponding probability of observing a specific realization. This formulation is particularly useful for the automated generation of large transition matrices with the aid of computer algebra systems, which we employed for the case of BTSP in the presence of correlated activity.

The standard method for understanding the dynamics of a Markov chain is to examine the eigenvalues and eigenvectors of its transition matrix. All matrices of this form always have one left eigenvector with equal entries and eigenvalue $\lambda_1 = 1$. The remaining eigenvalues have

$|\lambda_k| \leq 1$. In the simplest case, with a diagonalizable transition matrix and a single eigenvalue $= 1$, the time-evolution matrix can be written as

$$M^t = \sum_k \vec{r}_k \lambda_k^t \vec{l}_k \tag{3}$$

where $\vec{r}_k$ and $\vec{l}_k$ are the right- and left-eigenvectors of the transition matrix $M$ respectively, with normalization chosen so that $\vec{r}_k \cdot \vec{l}_k = 1$, and $\lambda_k$ are the eigenvalues, sorted in descending order of $|\lambda_k|$. As $t \to \infty$, any initial distribution over states described by the probability vector $\vec{x}(0)$ will converge to the unique stationary distribution corresponding to the probability vector $\vec{r}_1$. The asymptotic behavior is governed by the second leading eigenvalue:

$$\vec{x}(t) = M^t \vec{x}(0) = \sum_k \lambda_k^t \left(\vec{l}_k \cdot \vec{x}(0)\right) \vec{r}_k \sim \vec{r}_1 + \lambda_2^t \left(\vec{l}_2 \cdot \vec{x}(0)\right) \vec{r}_2 \tag{4}$$

where we have assumed that $\lambda_2$ has multiplicity $1$. In our case, the initial distribution is determined by the presentation of a particular pattern of activity to the neural network. Specifically, we characterize the relaxation dynamics to steady state of the subset of weights that are responsible for the encoding of that particular pattern, following the presentation of additional patterns. Assuming that the network has been operating under these dynamics for some time, the probability vector prior to exposure to this particular pattern will be described by $\vec{r}_1$. We provide examples of computing $\vec{x}(0)$ in the following sections.

#### Dicrete-time binary model of BTSP

In this section we define a BTSP plasticity rule for the case of binary weights, binary activity, and discrete time, examine the state space of the system, demonstrate how the learning can be transformed to a Markov chain, and quantify the scaling properties of this plasticity rule.

We discretize time into broad windows ($\approx 4s$) and introduce binary signals for the neurons, which differ depending on whether the neuron is pre- or post-synaptic for a given synapse. We first take into account the necessity of a postsynaptic plateau for plasticity, hence the postsynaptic signal is nonzero only if the time bin contained a plateau. From the presynaptic perspective, both place field activity and spikes accompanying a plateau potential contribute to the signal (SFig 10a). We then establish a rule for the potentiation and depotentiation of synapses, operating based solely on events taking place in the current and previous time bins: (i) if the synapse is depotentiated and there are both pre and postsynaptic signals in the same time bin, the synapse will be potentiated. (ii) conversely, if the synapse is potentiated and there is a postsynaptic signal at the current time bin with a presynaptic signal at the previous time bin (activity before plateau), or a postsynaptic signal at the previous time bin with a presynaptic signal at the current time bin (activity after plateau), then the synapse will be depotentiated (SFig 10b).

We then consider a synapse exposed to pre and postsynaptic signals $q_{pre}(t), q_{post}(t) \in \{0, 1\}$ and synaptic strength $W(t) \in \{0, 1\}$. The rule can then be written as

$$\begin{aligned} W(t+1) &= W(t) + (1 - W(t))\, q_{pre}(t) q_{post}(t) \\ &\quad - W(t)\, (q_{pre}(t) q_{post}(t-1) \vee q_{pre}(t-1) q_{post}(t)) \\ &= W(t) + (1 - W(t))\, q_{pre}(t) q_{post}(t) \\ &\quad - W(t) \Big( q_{pre}(t) q_{post}(t-1) + q_{pre}(t-1) q_{post}(t) \\ &\qquad - q_{pre}(t) q_{pre}(t-1) q_{post}(t) q_{post}(t-1) \Big) \end{aligned} \tag{5}$$

where the potentiation part of the rule is contained in the term with the $(1 - W(t))$ factor and the the depotentiation part of the rule is contained in the last term with the $-W(t)$ factor. Here we consider $q_{pre}$ and $q_{post}$ to be independent random variables sampled at each time step from Bernoulli distributions with probabilities $f_{pre}$ and $f_{post}$ respectively. These parameters define the "coding level" of the signals, where the smaller is the coding level, the sparser are the signals.

This process is non-Markovian due to the dependence on signals from time $t - 1$. In order to make this process Markovian, we introduce two additional variables, $\alpha(t) = W(t)q_{pre}(t-1)$ and $\beta(t) = W(t)q_{post}(t-1)$. We then derive the time evolution of $W$, $\alpha$, and $\beta$, found by multiplying Eq. 5 by 1, $q_{pre}(t)$, and $q_{post}(t)$, respectively, and using idempotence of $W$, $q_{pre}$, and $q_{post}$ (e.g., $W^2 = W$ due to $W$ being a binary variable) to derive (2). Now the dynamics of the variables $(W, \alpha, \beta)$ depend only on the current state of the system, making this is a Markov process. This process has five possible states, since $\alpha = 0$ and $\beta = 0$ if $W = 0$:

$$(W, \alpha, \beta) \in \{(0,0,0), (1,0,0), (1,1,0), (1,0,1), (1,1,1)\} \tag{6}$$

We can then derive the transition matrix of this process (Eq. 2):

$$M = \begin{pmatrix} 1-f_{pre}f_{post} & 0 & f_{post} & f_{pre} & 1-(1-f_{pre})(1-f_{post}) \\ 0 & (1-f_{pre})(1-f_{post}) & (1-f_{pre})(1-f_{post}) & (1-f_{pre})(1-f_{post}) & (1-f_{pre})(1-f_{post}) \\ 0 & f_{pre}(1-f_{post}) & f_{pre}(1-f_{post}) & 0 & 0 \\ 0 & (1-f_{pre})f_{post} & 0 & (1-f_{pre})f_{post} & 0 \\ f_{pre}f_{post} & f_{pre}f_{post} & 0 & 0 & 0 \end{pmatrix} \tag{7}$$

As described in the previous section, the transition matrix allows us to compute any probabilistic quantity related to the dynamics of an ensemble of independent synapses following the dynamics in Eq. 5. For instance, the steady-state probability of having a potentiated synapse (reported for $f_{pre} = f_{post} = f$ for simplicity):

$$P(W(\infty) = 1) = 1 - \vec{r}_1^{(1)} = \frac{1 - f + f^2}{3 - f - 2f^2 + 3f^3 - f^4} \tag{8}$$

where $\vec{r}_1^{(1)}$ denotes the probability of occupying the depotentiated $(0,0,0)$ state. All the remaining states are potentiated, so we can just subtract this quantity from 1 to obtain the steady state fraction of potentiated synapses.

We can also compute the second leading eigenvalue of this matrix (to leading order in the coding level of the pre and postsynaptic signals):

$$\lambda_2 \approx 1 - 3f_{pre}f_{post} + ... \tag{9}$$

which indicates an exponential approach to the steady state distribution with a time constant that scales with the product of the coding levels of the signals.

#### BTSP in neural networks

In order to further analyze the properties of this learning rule, we next consider a layer of $N_{pre}$ presynaptic neurons fully connected to a layer of $N_{post}$ postsynaptic neurons, with a weight matrix $W$. The analysis below also applies for the case of a recurrent network, where the input and output layers are assumed to be the same ($N_{pre} = N_{post} = N$). The weight matrix first learns associations between a stream of input patterns $\vec{\eta}_i(t) = q^{(i)}_{pre}(t), i = 1..N_{pre}$ and output patterns $\vec{\chi}_j(t) = q^{(j)}_{post}(t), j = 1..N_{post}$. After learning, we then examine the ability of this network to remember the learned associations. To this aim, we define the confusion matrix:

$$C(t_1, t_2) := \frac{\chi(t_1) \cdot (W\eta(t_2))}{N_{pre}N_{post}f_{pre}f_{post}} \tag{10}$$

which measures the similarity between the input currents $W\eta(t)$ to the output units and their "correct" activity level $\chi(t)$. This quantity will have a mean $C(\infty, \infty)$ for random patterns and will be measurably different for recently stored patterns. For this quantity, only the entries in the weight matrix corresponding to non-zero entries in the input and output vectors will contribute. Hence, we need to analyze the probability of a weight remaining in the potentiated state from the time it is exposed to a pattern with both high pre and postsynaptic signals. The subset of potentiated weights following the presentations of these patterns can be computed by considering the steady state probability vector $\vec{r}_1$ and operate on it with the transitions matrix $M$ (Eq. 7) where $f_{pre} = f_{post} = 1$.

$$\vec{x}(0) = \tilde{M}\vec{r}_1 = (\vec{r}_1^{(3)} + \vec{r}_1^{(4)} + \vec{r}_1^{(5)}, 0, 0, 0, \vec{r}_1^{(1)} + \vec{r}_1^{(2)}) \tag{11}$$

The time-evolution from this initial probability vector is then given by Eq. 4.

The quantity of interest is the signal-to-noise ratio (SNR), which can be derived from the confusion matrix $C$ (neglecting the correlations between synapses sharing the same presynaptic or postsynaptic neuron):

$$SNR(t) = \frac{(C(t,t) - C(\infty,\infty))^2}{var(C(t,t)) + var(C(\infty,\infty))} \tag{12}$$

where $C(\infty,\infty)$ represents random patterns obeying the same statistics as remembered patterns. The $SNR$ can be written as:

$$SNR(t) = N_{pre}N_{post}f_{pre}f_{post}\frac{(P(W(t)) - w)^2}{P(W(t))(1 - f_{pre}f_{post}P(W(t))) + w(1 - f_{pre}f_{post}w)} \tag{13}$$

where $P(W(t))$ is the probability that $W = 1$ at time $t$, and $w = \langle W \rangle$ is the steady state probability of having a potentiated synapse. The variance in the denominator arises from uncorrelated binomial distributions for the weights and pre/postsynaptic signals.

We can then estimate the SNR immediately following the presentation of a given pre and postsynaptic patterns (reported for simplicity for the case $f_{post} = f_{pre} = f$ and $N_{pre} = N_{post} = N$):

$$P(W(0)) = \frac{3 - 3f + f^2}{3 - f - 2f^2 + 3f^3 - f^4} \tag{14}$$

$$SNR(0) = N^2 f^2 \frac{(1-f)(3 - f - 2f^2 + 3f^3 - f^4)}{1 + 2f - 8f^2 + 13f^3 - 11f^4 + 5f^5 - f^6} \tag{15}$$

To derive an analytical expression for the number of retrievable patterns, we will assume that at threshold, the variance from the signal is of similar size to the variance of the noise. This results in

$$SNR(t) \sim \frac{N_{pre}N_{post}f_{pre}f_{post}}{2w(1 - f_{pre}f_{post}w)}\lambda_2^{2t} \tag{16}$$

The $SNR$ needs to scale with $\log(N)$ in order to successfully retrieve the output pattern in a recurrent network (*1, 2*). Hence, the maximum number of patterns that can be retrieved, $t^*$, is defined by $SNR(t^*) = s\log(N_{pre}N_{post})$, where $s$ is a proportionality constant chosen for the $SNR$ scaling, The memory capacity of the network is therefore:

$$t^* = -\frac{1}{2\log(\lambda_2)}\log\left(\frac{N_{pre}N_{post}f_{pre}f_{post}s\log(N_{pre}N_{post})}{2w(1 - f_{pre}f_{post}w)}\right) \tag{17}$$

We are most interested in sparse activity patterns, hence we can Taylor expand the second eigenvalue $\lambda_2$ in $f_{pre}$ and $f_{post}$, resulting in:

$$t^* \approx \frac{1}{6 f_{pre} f_{post}} \log\left(\frac{N_{pre} N_{post} f_{pre} f_{post} s \log(N_{pre} N_{post})}{2w(1 - f_{pre} f_{post} w)}\right) \tag{18}$$

With the scaling $f = \log(N)/N$, then

$$t^* \approx \frac{N_{pre} N_{post}}{6 \log(N_{pre}) log(N_{post})} \log\left(\frac{s \log(N_{pre} N_{post}) \log(N_{pre}) \log(N_{post})}{2w(1 - w)}\right) \tag{19}$$

Note that the $N^2$ scaling is optimal from an information theoretic perspective (*3–7*)

In SFig. 10e we compare the SNR from simulations of the BTSP rule (Eq. 5) with independent synapses and the prediction from the analysis based on the second leading eigenvalue (Eq. 16). The parameters used in Sfig. 10e are: $f_{post} = f$, $f_{pre} = 2f - f^2$ (to emulate a stream of presynaptic plateaus and activities, each with coding level $f$), $f = 10\frac{log(N)}{N}$, and $N = 1000$. The same parameters, but across different network sizes, were used in Fig. 5c for the capacity estimate based on the thresholded SNR from network simulations (where synapses are not independent). The threshold used was $s = 1$.

#### BTSP with correlated activity

We then investigated how this model would perform in the presence of temporal correlations in the stream of patterns, which are expected in realistic scenarios. These correlations might arise from correlations in the (processed) sensory inputs that any given neuron receives, or from the dynamics of a recurrent network, where the weights store information about previously encountered patterns. This realistic scenario is notoriously challenging for standard Hebbian rules, which only depend on the co-occurrence of pre and postsynaptic activity, due to the interference between similar patterns at storage and retrieval (Fig 5e, same-time rule curve).

Here we describe the full version of the model used for generating the results in Figure 5d and SFigure 10f,h. The difference, compared to the model in the previous section, is that we allow the activity component of the presynaptic signal to be correlated across time. We hence also need to account for the different presynaptic and postsynaptic plateaus that a synapse experience during learning. We modify Eq. 5 by substituting $q_{pre} \rightarrow a \vee q_{pre}$ where $q_{pre}$ denotes plateaus, and the activity $a$ is a two-state Markov process with transition probabilities $P(0 \rightarrow 1) = f_u = (1-c)f_a$ and $P(1 \rightarrow 0) = f_d = (1-c)(1-f_a)$, with $c$ denoting the temporal correlation of the activity, $< a_t a_{t+1} > - < a_t >^2 = c$, and $f_a$ the activity coding level (*8*).

In order to make the process Markovian, using the same definitions for $\alpha$ and $\beta$ as before, we include an additional variable $a_{-1}$ to represent the previous state of the activity. The resulting equations for the dynamics are:

$$\begin{cases} W' = W + (1-W)(a \vee q_{pre})q_{post} - W(a \vee q_{pre})\beta \\ \qquad - W(a_{-1} + \alpha - a_{-1}\alpha)q_{post}\left(1 - \beta(a \vee q_{pre})\right), \\ \alpha' = Wq_{pre} + (1-W)(a \vee q_{pre})q_{pre}q_{post} - W(a \vee q_{pre})q_{pre}\beta \\ \qquad - W(a_{-1} + \alpha - a_{-1}\alpha)q_{pre}q_{post}\left(1 - \beta(a \vee q_{pre})\right), \\ \beta' = Wq_{post} + (1-W)(a \vee q_{pre})q_{post} - W(a \vee q_{pre})q_{post}\beta \\ \qquad - W(a_{-1} + \alpha - a_{-1}\alpha)q_{post}\left(1 - \beta(a \vee q_{pre})\right), \\ a' = a_{-1} + (1-a_{-1})q_u - a_{-1}q_d \end{cases} \tag{20}$$

where $q_u$ and $q_d$ are Bernoulli variables with probabilities $f_u$ and $f_d$, respectively, and $a_{-1}$, $q_{pre}$, $q_{post}$, and $W$ are all idempotent. As done above, we can examine the transition matrix associated to this process. The available states are

$$(W, \alpha, \beta, a_{-1}) \in \Big\{(0,0,0,0), (0,0,0,1), (1,0,0,0), (1,1,0,0), (1,0,1,0), \\ (1,0,0,1), (1,1,1,0), (1,1,0,1), (1,0,1,1), (1,1,1,1)\Big\} \tag{21}$$

with a transition probability matrix (for $f_{pre} = f_{post} = f_p$)

$$
\begin{pmatrix}
\left(1-f_p^2\right)(1-(1-c)f_a) & (1-c)(1-f_a)\left(1-f_p^2\right) & 0 & f_p(1-(1-c)f_a) & f_p(1-(1-c)f_a) & f_p(1-(1-c)f_a) & (1-c)(1-f_a)f_p & (1-c)(1-f_a)\left(2f_p-f_p^2\right) & (1-c)(1-f_a)\left(2f_p-f_p^2\right) & (1-c)(1-f_a)\left(2f_p-f_p^2\right) \\
(1-f_p)(1-c)f_a & (1-(1-c)(1-f_a))(1-f_p) & 0 & f_p(1-c)f_a & (1-c)f_a & (1-(1-c)(1-f_a))f_p & (1-c)f_a & 1-(1-c)(1-f_a) & 1-(1-c)(1-f_a) & 1-(1-c)(1-f_a) \\
0 & 0 & (1-f_p)^2(1-(1-c)f_a) & (1-f_p)^2(1-(1-c)f_a) & (1-f_p)^2(1-(1-c)f_a) & (1-c)(1-f_a)(1-f_p)^2 & (1-f_p)^2(1-(1-c)f_a) & (1-c)(1-f_a)(1-f_p)^2 & (1-c)(1-f_a)(1-f_p)^2 & (1-c)(1-f_a)(1-f_p)^2 \\
0 & 0 & (1-f_p)f_p(1-(1-c)f_a) & (1-f_p)f_p(1-(1-c)f_a) & 0 & (1-c)(1-f_a)(1-f_p)f_p & 0 & (1-c)(1-f_a)(1-f_p)f_p & 0 & 0 \\
0 & 0 & (1-f_p)f_p(1-(1-c)f_a) & 0 & (1-f_p)f_p(1-(1-c)f_a) & 0 & 0 & 0 & 0 & 0 \\
0 & 0 & (1-f_p)^2(1-c)f_a & (1-f_p)^2(1-c)f_a & 0 & (1-(1-c)(1-f_a))(1-f_p)^2 & 0 & (1-(1-c)(1-f_a))(1-f_p)^2 & 0 & 0 \\
f_p^2(1-(1-c)f_a) & (1-c)(1-f_a)f_p^2 & f_p^2(1-(1-c)f_a) & 0 & 0 & 0 & 0 & 0 & 0 & 0 \\
0 & 0 & (1-f_p)f_p(1-c)f_a & (1-f_p)f_p(1-c)f_a & 0 & (1-(1-c)(1-f_a))(1-f_p)f_p & 0 & (1-(1-c)(1-f_a))(1-f_p)f_p & 0 & 0 \\
(1-f_p)f_p(1-c)f_a & (1-(1-c)(1-f_a))(1-f_p)f_p & (1-f_p)f_p(1-c)f_a & 0 & 0 & 0 & 0 & 0 & 0 & 0 \\
f_p^2(1-c)f_a & (1-(1-c)(1-f_a))f_p^2 & f_p^2(1-c)f_a & 0 & 0 & 0 & 0 & 0 & 0 & 0
\end{pmatrix} \tag{22}
$$

From this, the eigenvalues can be calculated, and the second leading eigenvalue is

$$
\lambda_2 \sim 1 + \left((-3 + c + c^2)f_a - 3f_p\right) f_p \tag{23}
$$

Note that, in the presence of correlations, the quadratic scaling of this eigenvalue with the square of the coding level still holds. Additionally, increasing the activity correlation $c$ brings $\lambda_2$ closer to $1$, resulting in an increased capacity. The depotentiation component of the rule plays a crucial role in ensuring the network's robustness to temporal correlations in the activity patterns. As the depotentiation process is triggered when there is a presynaptic signal followed by a postsynaptic signal or vice versa, it effectively decorrelates the signal across neighboring time bins. This means that if a specific pattern of activity is correlated with the previous pattern, the depotentiation process helps to prevent the current pattern from interfering with or being influenced by the previously stored pattern. In the presence of temporal correlations, depotentiation events become targeted rather than random, which allows BTSP to store more attractors. This targeted depotentiation aids in reducing interference between similar patterns and enhances the network's ability to store and retrieve information in realistic scenarios with correlated inputs (Sfig. 11).

In order to analyze the dynamics of this chain, one must define the initial probability vector $\vec{x}(0)$. Here, we we might be interested in three different versions of $SNR$, corresponding to different selections of the subset of synaptic weights: (i) only the units which had a presynaptic plateau and postsynaptic plateau ($\vec{x}_{(p)}(0)$) (ii) presynaptic activity and postsynaptic plateau ($\vec{x}_{(a)}(0)$) (iii) presynaptic plateau or activity and postsynaptic plateau ($\vec{x}_{(a\vee p)}(0)$). Given the steady-state probability vector $\vec{r}_1^{(i)}$, we can calculate the entries in $\vec{x}(0)$. For instance, in $\vec{x}_{(a)}(0)$, we are projecting out states with $a_{-1} \neq 0$. Steady state transitions from the state $(0,0,0,0)$ to $(1,1,1,0)$ happen with probability $(1-f_u)f_p^2$, so we have a contribution from $\vec{r}_1^{(1)}$ with this coefficient. Similar considerations lead to:

$$\vec{x}_{(p)}(0) = \frac{f_p^2}{Z_{(p)}} \begin{pmatrix} f_d(\vec{r}_1^{(6)}+\vec{r}_1^{(8)}+\vec{r}_1^{(9)}+\vec{r}_1^{(10)})+(1-f_u)(\vec{r}_1^{(4)}+\vec{r}_1^{(5)}+\vec{r}_1^{(7)}) \\ (1-f_d)(\vec{r}_1^{(6)}+\vec{r}_1^{(8)}+\vec{r}_1^{(9)}+\vec{r}_1^{(10)})+f_u(\vec{r}_1^{(4)}+\vec{r}_1^{(5)}+\vec{r}_1^{(7)}) \\ 0 \\ 0 \\ 0 \\ 0 \\ (1-f_u)(\vec{r}_1^{(1)}+\vec{r}_1^{(3)})+f_d(\vec{r}_1^{(2)}) \\ 0 \\ 0 \\ f_u(\vec{r}_1^{(1)}+\vec{r}_1^{(3)})+(1-f_d)\vec{r}_1^{(2)} \end{pmatrix} \tag{24}$$

$$\vec{x}_{(a)}(0) = \frac{f_p}{Z_{(a)}} \begin{pmatrix} 0 \\ (1-f_d)(\vec{r}_1^{(6)}+\vec{r}_1^{(8)}+\vec{r}_1^{(9)}+\vec{r}_1^{(10)})+f_u(\vec{r}_1^{(4)}+\vec{r}_1^{(5)}+\vec{r}_1^{(7)}) \\ 0 \\ 0 \\ 0 \\ 0 \\ 0 \\ 0 \\ (1-f_p)\left(f_u(\vec{r}_1^{(1)}+\vec{r}_1^{(3)})+(1-f_d)\vec{r}_1^{(2)}\right) \\ f_p\left(f_u(\vec{r}_1^{(1)}+\vec{r}_1^{(3)})+(1-f_d)\vec{r}_1^{(2)}\right) \end{pmatrix} \tag{25}$$

$$\vec{x}_{(a \vee p)}(0) = \frac{f_p}{Z_{(a \vee p)}} \begin{pmatrix} f_p\Big(f_d(\vec{r}_1^{(6)}+\vec{r}_1^{(8)}+\vec{r}_1^{(9)}+\vec{r}_1^{(10)})+(1-f_u)(\vec{r}_1^{(4)}+f_p\vec{r}_1^{(5)}+\vec{r}_1^{(7)})\Big) \\ (1-f_d)(\vec{r}_1^{(6)}+\vec{r}_1^{(8)}+\vec{r}_1^{(9)}+\vec{r}_1^{(10)})+f_u(\vec{r}_1^{(4)}+\vec{r}_1^{(5)}+\vec{r}_1^{(7)}) \\ 0 \\ 0 \\ 0 \\ 0 \\ f_p\Big((1-f_u)(\vec{r}_1^{(1)}+\vec{r}_1^{(3)})+f_d(\vec{r}_1^{(2)})\Big) \\ 0 \\ (1-f_p)\Big(f_u(\vec{r}_1^{(1)}+\vec{r}_1^{(3)})+(1-f_d)\vec{r}_1^{(2)}\Big) \\ f_p\Big(f_u(\vec{r}_1^{(1)}+\vec{r}_1^{(3)})+(1-f_d)\vec{r}_1^{(2)}\Big) \end{pmatrix} \tag{26}$$

where the normalization constants $Z$, which enter in the scaling factor for the SNR, are the fraction of synapses participating in the encoding:

$$Z_{(p)} = f_p^2 \tag{27}$$

$$Z_{(a)} = f_a f_p \tag{28}$$

$$Z_{(a \vee p)} = f_p(f_p + f_a - f_p f_a) \tag{29}$$

We can then proceed with computing the probability of $W = 1$ using the methods above (note that now both of the first two dimensions contain $W = 0$). We will present the results from $\vec{x}_{(a \vee p)}(0)$ in SFig. 10f, comparing the SNR from simulations of the BTSP rule in the presence of correlations (Eq. 20) with independent synapses and the prediction from the analysis based on the full theory as well as the approximation using the second leading eigenvalue (Eq. 23). The parameters used in Sfig. 10f are: $f_p = 10\log(N)/N$, $f_a = 5\log(N)/N$, $c = .8$ (to emulate a stream of correlated activity in a network paired with a similar number of plateaus), and $N = 800$. In Fig. 5d, we used $f_p = f_a = 10\log(N)/N$ and $N = 3000$ as we varied $c$.

We show the confusion matrix (SFig. 10h) and attractor capacity (Fig. 5d, see section "Network dynamics" in Methods) from network simulations using the BTSP rule in the presence of correlations, with parameters: $N = 8000$, $f_p = 5\frac{log(N)}{N}$, $f_a = 10\frac{log(N)}{N}$, averaged over 30 repetitions. The correlation used for Sfig. 10h was $c = 0.8$.

#### Same-time rule

We modified the Hebbian plasticity rule, explored in depth in [Fusi 94], to include correlated activity. A synapse is potentiated if pre and postsynaptic activity are coincident in time and stochastically depotentiated (with probability $f_{dep}$) whenever pre-synaptic activity coincides with post-synaptic silence (or viceversa). The synaptic dynamics for this rule in the presence of correlations are:

$$\begin{cases} W' = W + (1-W)(q_{pre} \vee a_{pre})(q_{post} \vee a_{post}) \\ \qquad - W\left((q_{pre} \vee a_{pre})(1 - q_{post} \vee a_{post}) + (1 - q_{pre} \vee a_{pre})(q_{post} \vee a_{post})\right) q_{dep} \\ a'_{pre} = a_{pre} + (1 - a_{pre})q_{u,pre} - a_{pre}q_{d,pre} \\ a'_{post} = a_{post} + (1 - a_{post})q_{u,post} - a_{post}q_{d,post} \end{cases} \tag{30}$$

where $q_{dep}$ is sampled from a Bernoulli distribution with parameter $f_{dep}$.

As in the BTSP rule, the presynaptic signal contains both a correlated component (which was due to activity in BTSP) and an uncorrelated component (which was due to plateaus in BTSP). Hence, the patterns statistics are matched to those used for BTSP. Since, in the standard Hebbian rule, there is no distinction between plateaus and activity, the postsynaptic signal includes both for this rule. We estimated confusion matrices and attractor capacity (see next section) based on the uncorrelated component of the output signal, since the post-synaptic signal for BTSP is determined by the uncorrelated plateaus. The confusion matrix (Sfig. 10g) and attractor capacity (Fig 5d) are from network simulations with parameters identical to those used in BTSP (see previous section).

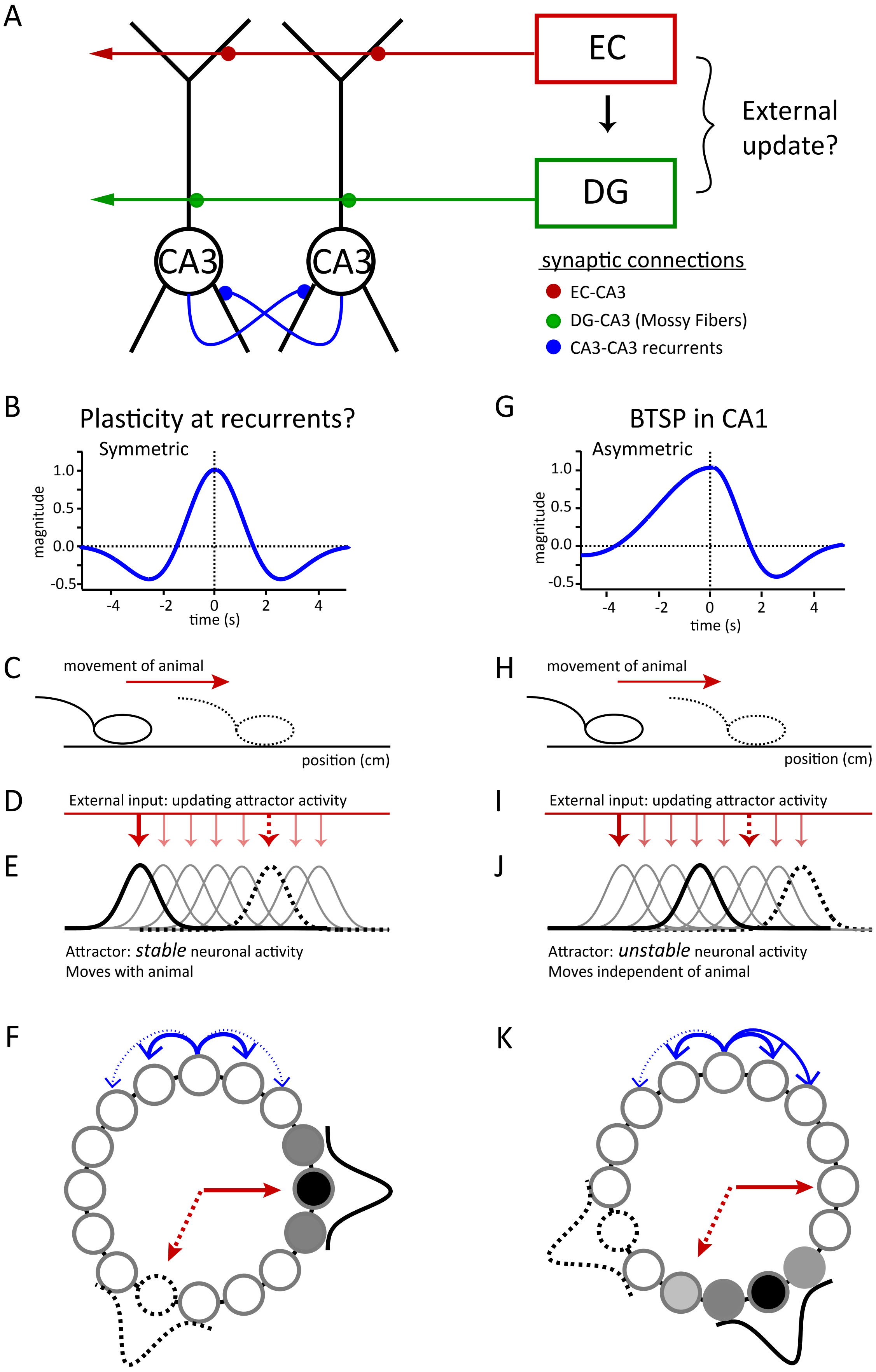
A
EC
DG
CA3
CA3
External update?
synaptic connections
EC-CA3
DG-CA3 (Mossy Fibers)
CA3-CA3 recurrents
B
Plasticity at recurrents?
Symmetric
magnitude
1.0
0.5
0.0
-0.5
-4
-2
0
2
4
time (s)
G
BTSP in CA1
Asymmetric
magnitude
1.0
0.5
0.0
-0.5
-4
-2
0
2
4
time (s)
C
movement of animal
position (cm)
H
movement of animal
position (cm)
D
External input: updating attractor activity
E
Attractor: *stable* neuronal activity
Moves with animal
I
External input: updating attractor activity
J
Attractor: *unstable* neuronal activity
Moves independent of animal
F
K

**Fig. S1. Possible attractor formation mechanisms in CA3.** (**A**) CA3 circuit schematic showing various excitatory synaptic input pathways and their approximate locations on a pair of CA3 pyramidal neurons compromising a single attractor. (**B**) A bidirectional synaptic plasticity rule expressing a symmetric time course of potentiation (positive values) and de-potentiation (negative values). Such a plasticity rule applied to recurrent synapses will produce a stable activity pattern that persist at a given location. To link the updating of this activity pattern with the (**C**) behavior of an animal (i.e. produce a different set of neurons active at different locations) (solid figure is current location; dashed figure is future location) (**D**) an additional excitatory input is needed to update the activity in the attractor network based on the movement of the animal. (**E**) (solid dark red arrows are current active external input, solid black gaussian is current activity, dark dashed arrows and gaussian are signals at future active locations and light solid arrows and gaussians are signals at intervening active locations). (**F**) A ring version[57] of this attractor system where a symmetric rule (blue lines; solid is potentiation and dashed depotentiation) combined with uniform inhibition (not shown) produces a bump of activity (black gaussian at black circles) whose movement is determined by an updating mechanism linked to behavior (red arrows). Shading of circles around bump indicates symmetric spread of synaptic current. (**G**) BTSP in CA1 has an asymmetric time-course that will produce activity pattern trajectories that are independent of animal behavior (**H** to **K**) (same symbolic structure as C to F). Shading of circles around bump indicates asymmetric spread of synaptic current. The asymmetry present in CA1 BTSP has been hypothesized to result from activity traces with different time courses (ETs > PTs). Symmetric rules can be achieved when the activity traces are equal in duration (ET=PT). Activity traces are thought to be biochemical filters of synaptic input (ETs) and plateau potentials (PTs)[25,27,40].

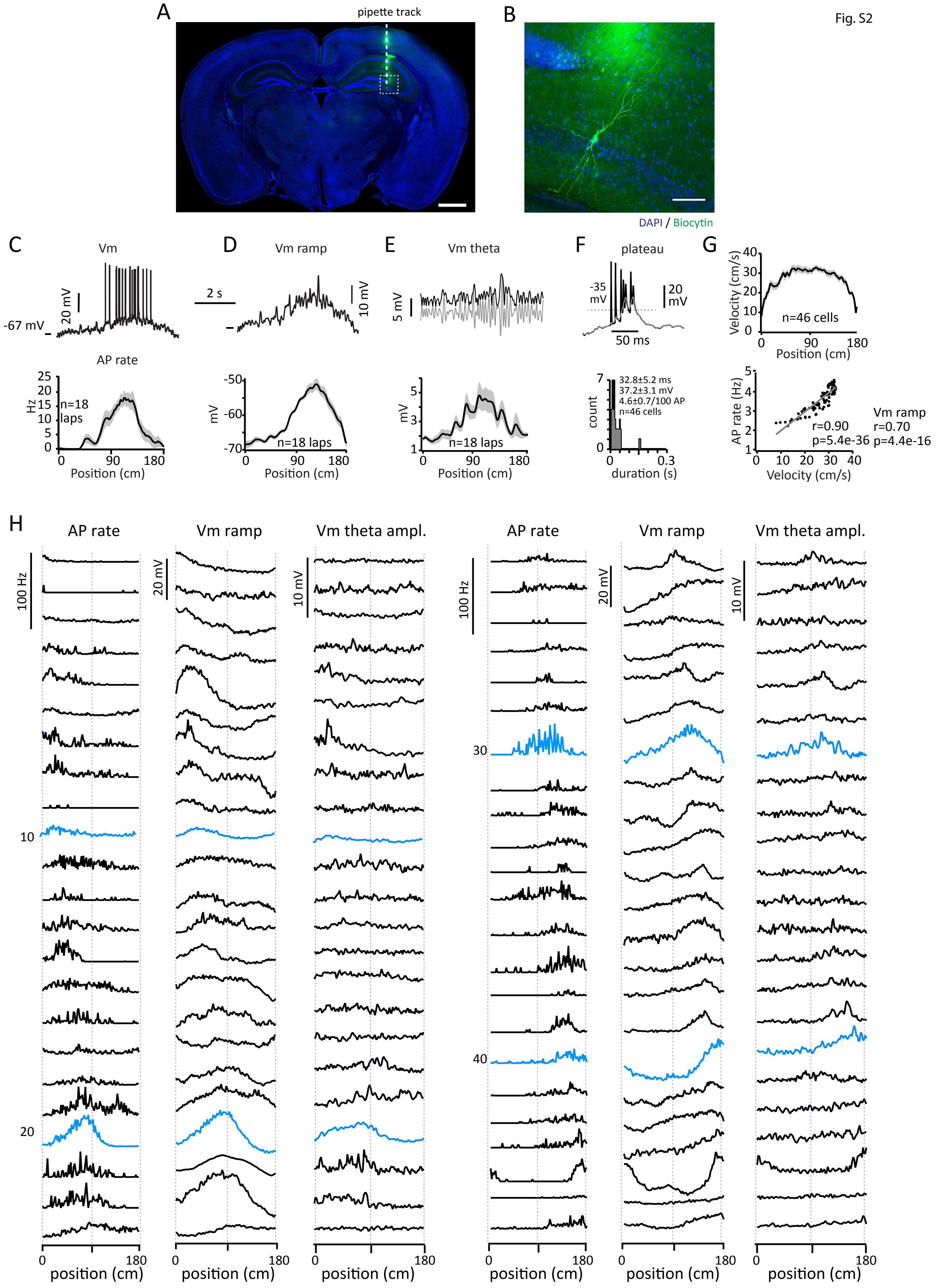

A
pipette track
B
DAPI / Biocytin
C
Vm
-67 mV
20 mV
AP rate
Hz
n=18 laps
Position (cm)
D
Vm ramp
2 s
10 mV
mV
n=18 laps
Position (cm)
E
Vm theta
5 mV
mV
n=18 laps
Position (cm)
F
plateau
-35 mV
20 mV
50 ms
count
32.8±5.2 ms
37.2±3.1 mV
4.6±0.7/100 AP
n=46 cells
duration (s)
G
Velocity (cm/s)
n=46 cells
Position (cm)
AP rate (Hz)
r=0.90
p=5.4e-36
Vm ramp
r=0.70
p=4.4e-16
Velocity (cm/s)
H
AP rate
Vm ramp
Vm theta ampl.
100 Hz
20 mV
10 mV
10
20
30
40
position (cm)

**Fig. S2. Whole-cell recordings from hippocampal area CA3.** (**A**) Histology shows electrode track for recording targeting CA3, scale bar 1 mm. (**B**) Enlarged image shows the recorded CA3 pyramidal neuron filled with biocytin, scale bar 100 um. (**C**) Vm, (**D**) Vm ramp and (**E**) Vm theta expanded from lap3 (upper) and average spatial profile for multiple laps (lower) for cell shown in Fig. 1. (**F**) Vm expanded from lap 5 showing plateau associated burst firing with threshold for plateau shown (dashed line). Histogram of plateau duration for all PCs (lower). (**G**) (upper) Average spatial profile for mouse run velocity from all mice during PC recordings. (lower) PC AP firing rate is strongly correlated with mouse running velocity. Values for Vm/velocity correlation also listed. (**H**) Spatially binned (100 bins of 1.8 cm) unsmoothed average AP rate, Vm ramp, and Vm theta amplitude (averages from 5-48 trials) for all 46 PCs recorded from CA3. Recordings are sorted by peak AP rate location.

Fig. S3

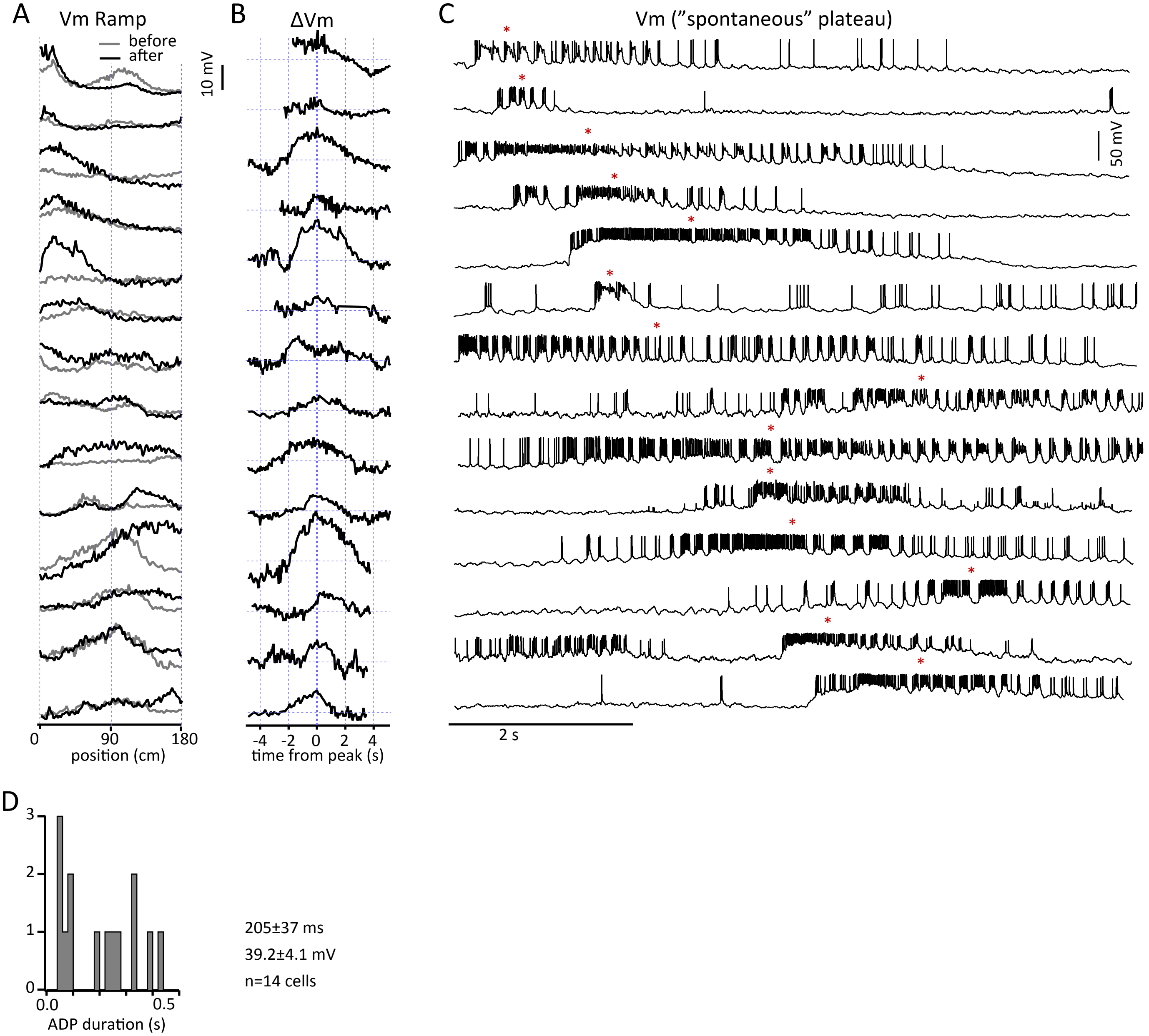

A
Vm Ramp
before
after
10 mV
0
90
180
position (cm)
B
ΔVm
-4
-2
0
2
4
time from peak (s)
C
Vm ("spontaneous" plateau)
50 mV
2 s
D
3
2
1
0
0.0
0.5
ADP duration (s)
205±37 ms
39.2±4.1 mV
n=14 cells

**Fig. S3. Naturally occurring long-duration plateaus in CA3 neurons.** (**A**) Vm ramps (gray; before plateaus, black; after plateaus) (**B**) and ΔVm from all 14 neurons exhibiting (**C**) naturally occurring long-duration plateau potentials. Red asterisks denote location (time) of peak positive ΔVm. (**D**) Histogram of ADP duration during naturally occurring plateau potentials.

Fig. S4

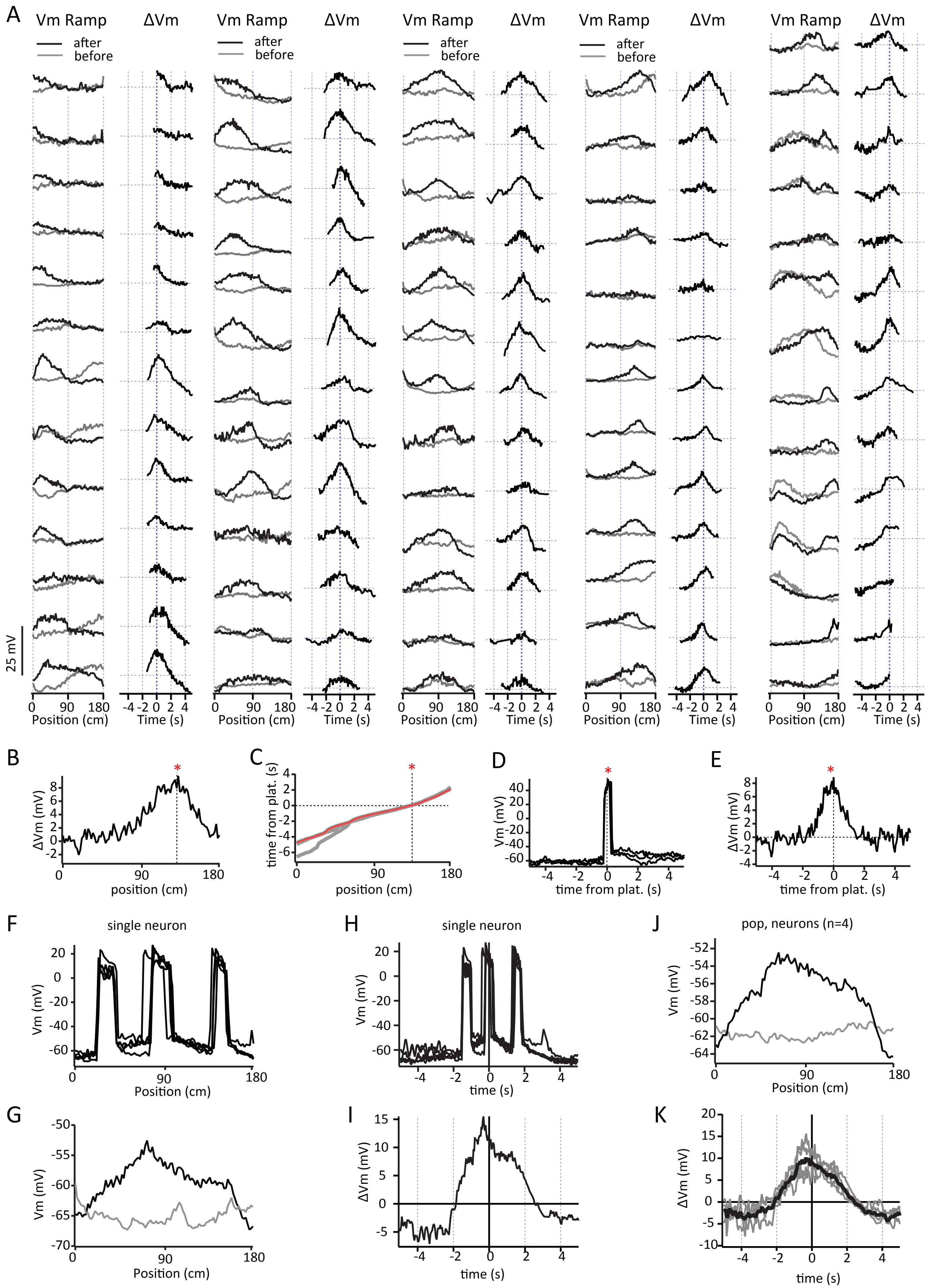

A
Vm Ramp
ΔVm
after
before
25 mV
Position (cm)
Time (s)
B
ΔVm (mV)
position (cm)
C
time from plat. (s)
position (cm)
D
Vm (mV)
time from plat. (s)
E
ΔVm (mV)
time from plat. (s)
F
single neuron
Vm (mV)
Position (cm)
H
single neuron
Vm (mV)
time (s)
J
pop, neurons (n=4)
Vm (mV)
Position (cm)
G
Vm (mV)
Position (cm)
I
ΔVm (mV)
time (s)
K
ΔVm (mV)
time (s)

**Fig. S4. Induced plateaus generate PFs in CA3 neurons.** (**A**) Spatially binned Vm ramp (gray, before induction, black; after induction) from all 66 recordings. Change in Vm versus time from plateau (black, ΔVm). (**B**) Spatially binned change in Vm ramp caused by three trials of induced plateau plotted in space. (**C**) Spatially binned time from the location of the plateau induction for three induction trials (gray). Red trace is briefest time from plateau initiation site for the three induction trials. This red trace is used as time base for this cell. (**D**) Three spatially binned induction trials (Vm with APs removed) plotted versus the time base produced above showing the location in time of current injections. (**E**) Spatially binned change in Vm ramp plotted versus time base. Same cell as shown in Fig. 2. Red asterisk in (B to E) indicate location of plateau induction. (**F**) Five spatially binned induction trials (Vm with APs removed) showing induction currents that were injected at three different locations across the distance of the track within the same trials in a single neuron (~30, 90, 145 cm). (**G**) Spatially binned Vm ramps (gray; before induction, black; after induction) from same neuron. (**H**) The same five spatially binned induction trials now plotted versus time base showing that they are centered on the middle current injection. (**I**) Spatially binned change in Vm ramp caused by five trials of induced plateau plotted in time in the same single neuron. (**J**) Average spatially binned Vm ramps (gray; before inductions, black; after inductions) from population of four neurons receiving the same multiple induction protocol. (**K**) Spatially binned change in Vm ramp caused by multiple induction protocol plotted in time for four neurons (gray; individual neurons, black; average). Note that the time course of the change is similar to that induced by a single current injection (fig. S5D). Induction protocols in (F to K) were used to mimic trains of plateaus.

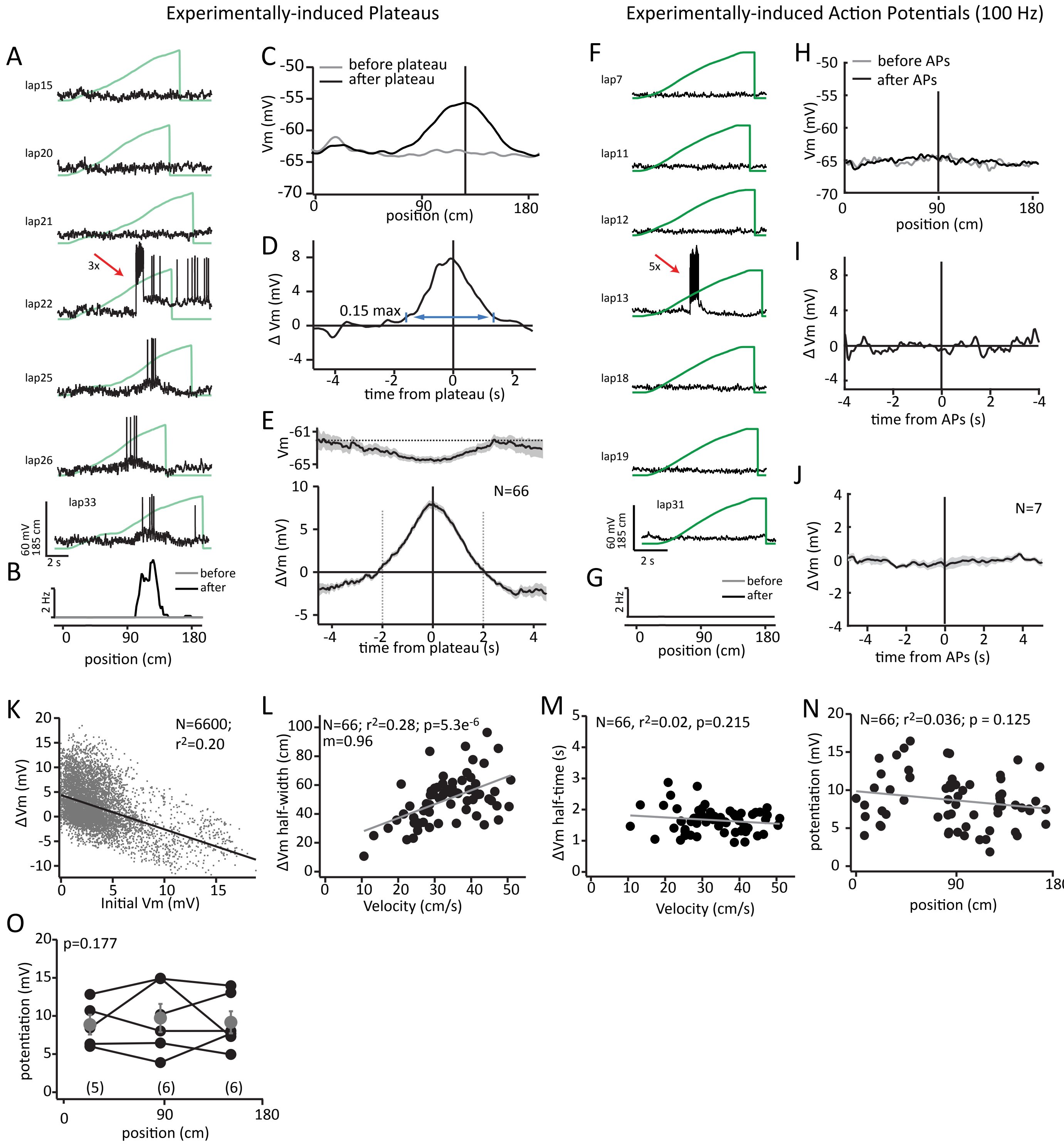

Experimentally-induced Plateaus
Experimentally-induced Action Potentials (100 Hz)
A
lap15
lap20
lap21
3x
lap22
lap25
lap26
lap33
60 mV
185 cm
2 s
B
before
after
2 Hz
position (cm)
C
before plateau
after plateau
Vm (mV)
position (cm)
D
Δ Vm (mV)
0.15 max
time from plateau (s)
E
Vm
N=66
ΔVm (mV)
time from plateau (s)
F
lap7
lap11
lap12
5x
lap13
lap18
lap19
lap31
60 mV
185 cm
2 s
G
before
after
2 Hz
position (cm)
H
before APs
after APs
Vm (mV)
position (cm)
I
Δ Vm (mV)
time from APs (s)
J
N=7
Δ Vm (mV)
time from APs (s)
K
N=6600;
r2=0.20
ΔVm (mV)
Initial Vm (mV)
L
N=66; r2=0.28; p=5.3e-6
m=0.96
ΔVm half-width (cm)
Velocity (cm/s)
M
N=66, r2=0.02, p=0.215
ΔVm half-time (s)
Velocity (cm/s)
N
N=66; r2=0.036; p = 0.125
potentiation (mV)
position (cm)
O
p=0.177
potentiation (mV)
(5)
(6)
(6)
position (cm)

**Fig. S5. Additional properties of CA3 BTSP.** (**A**) Membrane potential (Vm; black) and mouse position (green) for laps before (15-21) during (22) and after (25-26, 33) plateau induction. Average firing rate (**B**) and Vm ramp (**C**) for laps before (gray) and after (black) plateau induction. (**D**) Difference between Vm ramp after and before plateau (ΔVm) from PC in (A). Arrow shows region of Vm ramp where duration calculations were performed. (**E**) Average Vm for trials before induction (upper) and ΔVm (lower) for population of neurons with induced plateaus. (**F**) Membrane potential (Vm; black) and mouse position (green) for laps before (7, 11 and 12) during (13) and after (18, 19 and 31) 100 Hz spike train induction (5 laps total). Average firing rate (**G**) average Vm ramp (**H**) for laps before (gray) and after (black) 100 Hz spike train induction. (**I**) Difference between Vm ramp after and before 100 Hz spike train induction (ΔVm) from PC in (F). (**J**) Average ΔVm for population of neurons after 100 Hz spike train induction. (**K**) Dependence of bidirectional Vm change on initial Vm. (**L**) Relationship of Vm ramp width in space and velocity of mouse during induction trials. (**M**) Relationship of Vm ramp width in time and velocity of mouse during induction trials. (**N**) Relationship between the amplitude of BTSP potentiation and position of induction for the population. (**O**) Relationship between the amplitude of BTSP potentiation and position of induction for individual neurons with multiple inductions at different locations.

Fig. S6

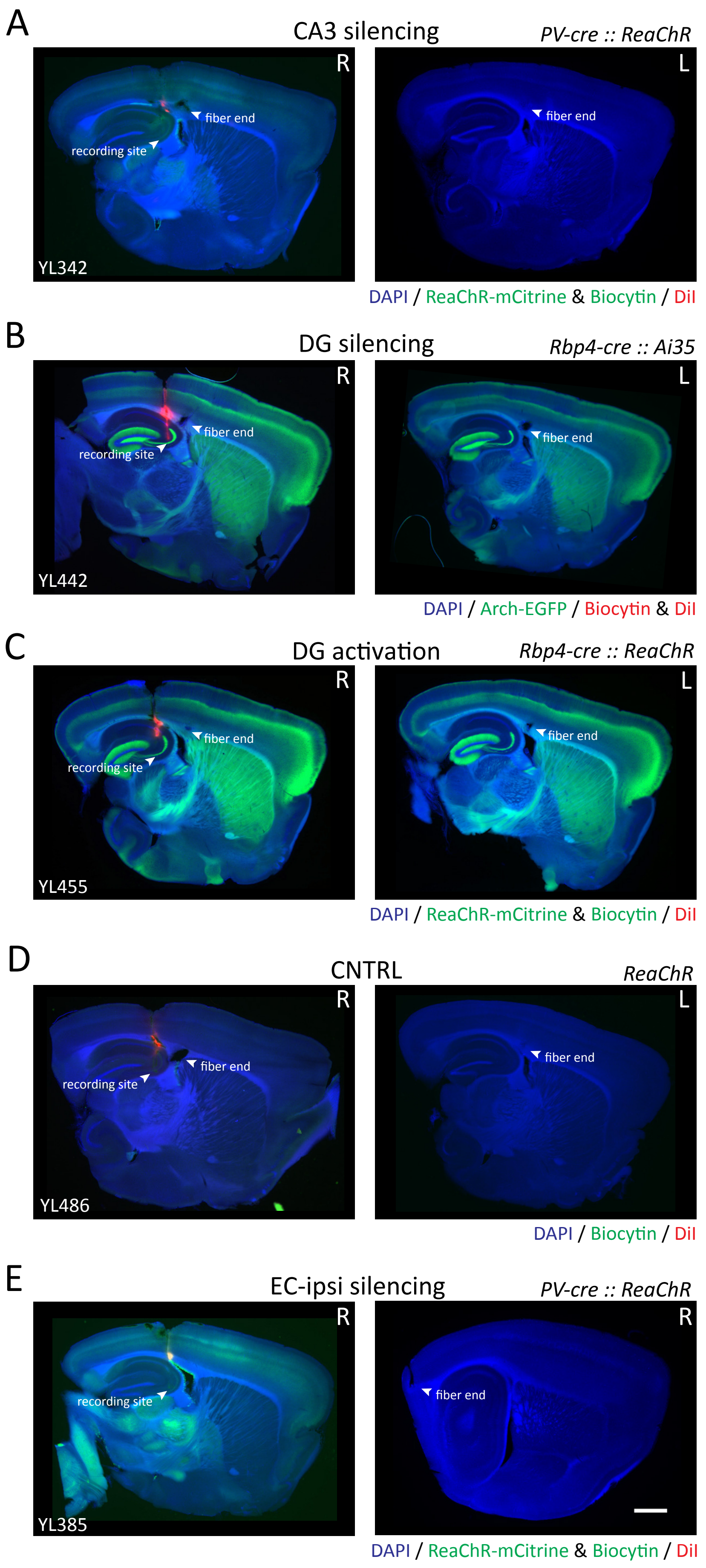

A
CA3 silencing
PV-cre :: ReaChR
R
L
fiber end
recording site
fiber end
YL342
DAPI / ReaChR-mCitrine & Biocytin / DiI
B
DG silencing
Rbp4-cre :: Ai35
R
L
fiber end
recording site
fiber end
YL442
DAPI / Arch-EGFP / Biocytin & DiI
C
DG activation
Rbp4-cre :: ReaChR
R
L
fiber end
recording site
fiber end
YL455
DAPI / ReaChR-mCitrine & Biocytin / DiI
D
CNTRL
ReaChR
R
L
fiber end
recording site
fiber end
YL486
DAPI / Biocytin / DiI
E
EC-ipsi silencing
PV-cre :: ReaChR
R
R
recording site
fiber end
YL385
DAPI / ReaChR-mCitrine & Biocytin / DiI

**Fig. S6. Expression of optogenetic actuators and optical fiber implantation.** Histology shows the optical fiber and pipette tracks for (**A**) CA3 silencing, (**B**) control, (**C**) DG silencing, (**D**) DG activation and (**D**) ipsi-EC silencing. Arrows indicate the recording site in CA3 and optical fiber end on the hippocampal CA3 or EC. R and L indicate right and left hemisphere, respectively. Scale bar 1 mm.

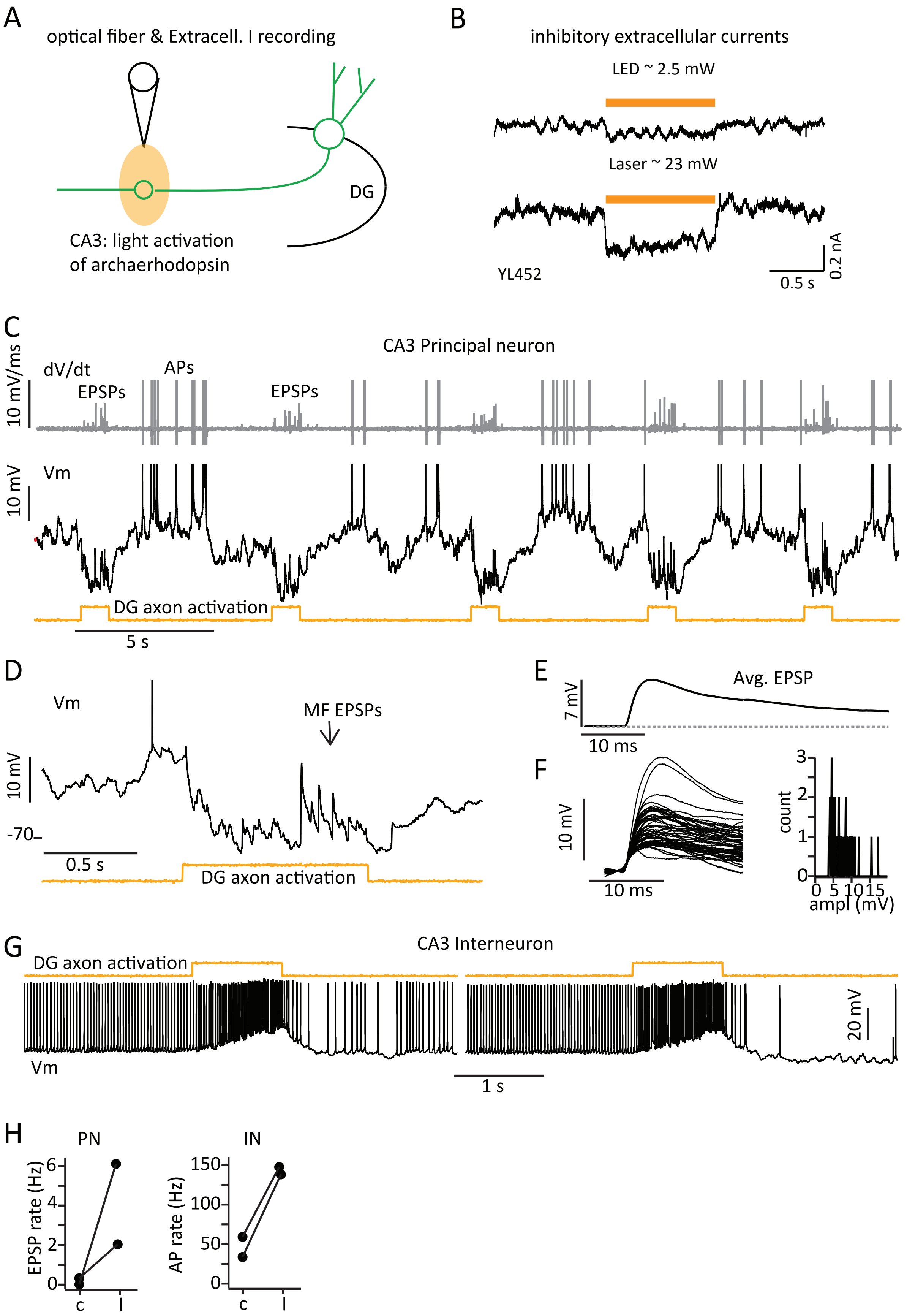
A
optical fiber & Extracell. I recording
DG
CA3: light activation
of archaerhodopsin
B
inhibitory extracellular currents
LED ~ 2.5 mW
Laser ~ 23 mW
YL452
0.5 s
0.2 nA
C
CA3 Principal neuron
10 mV/ms
dV/dt
EPSPs
APs
EPSPs
10 mV
Vm
DG axon activation
5 s
D
Vm
MF EPSPs
10 mV
-70
0.5 s
DG axon activation
E
7 mV
Avg. EPSP
10 ms
F
10 mV
10 ms
count
ampl (mV)
G
CA3 Interneuron
DG axon activation
20 mV
Vm
1 s
H
PN
IN
EPSP rate (Hz)
AP rate (Hz)
c
l

**Fig. S7. Confirmation of DG manipulation.** (**A**) Sketch of manipulation condition. (**B**) Optical activation (yellow) of Archaerhodopsin in DG produces large amplitude, hyperpolarization related local field currents (sources) in the CA3. (**C**) Optical activation of ReaChR expression in DG (yellow, lower) drives activity in CA3 principal neuron. Vm (black, middle) and temporal derivative (gray; upper) show fast rising large amplitude EPSP (2-20 mV/V) and AP related transients (>20 mV/V). (**D**) Expansion of Vm for last light activation. (**E**) Avg. EPSP events, aligned at dV/dt threshold (2 mV/ms), from 10 light activation periods. (**F**) Individual EPSP events and histogram of event amplitudes. (**G**) Optical activation of ReaChR expression in DG (yellow, upper) drives activity in CA3 fast-spiking interneuron. Vm (black, lower) traces for two separate activations shows increase in firing rate. (**H**) Increases in large amplitude, fast rising EPSP rates in principal neurons (n=2) and AP rates in 2 interneurons.

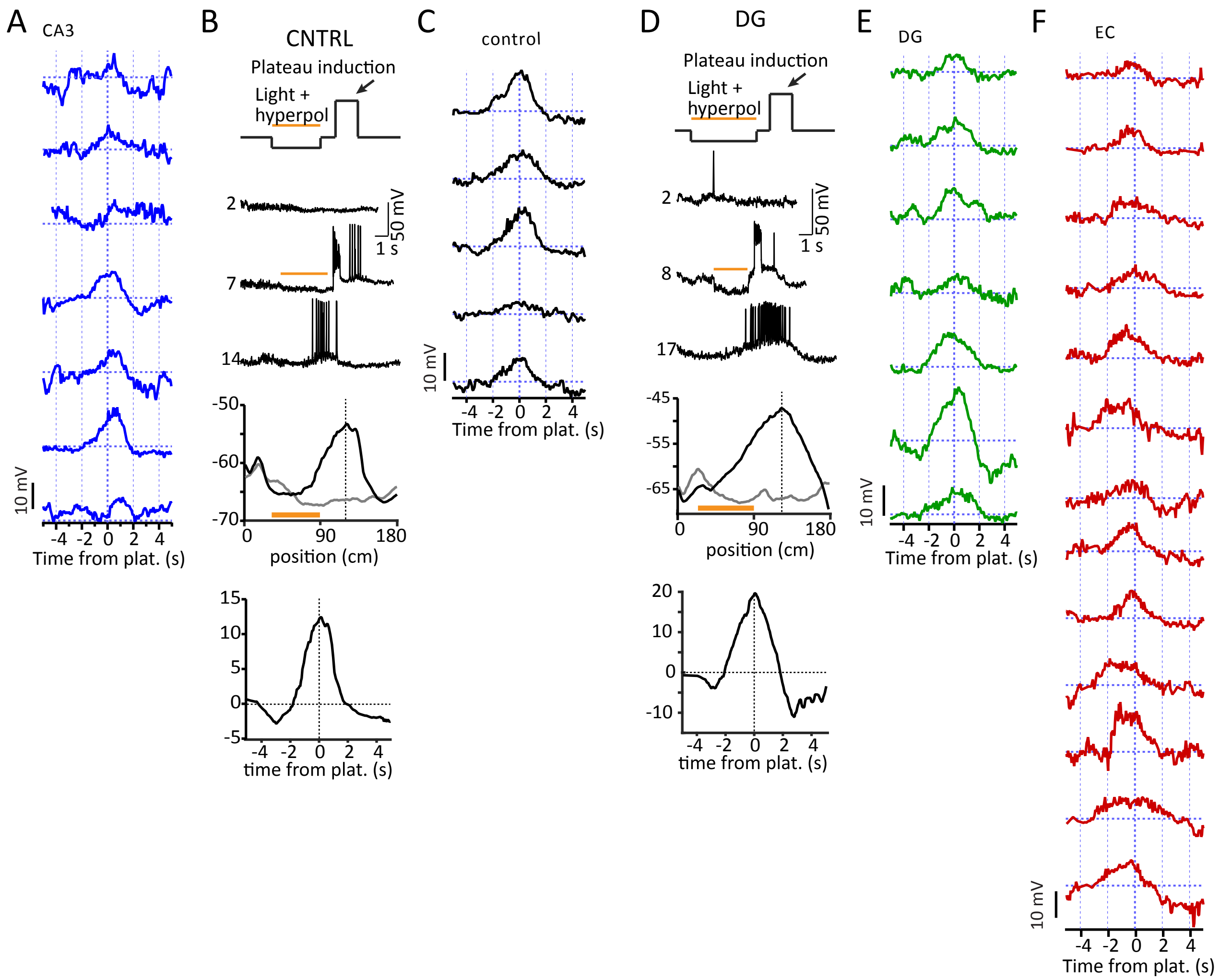
A
CA3
10 mV
-4 -2 0 2 4
Time from plat. (s)
B
CNTRL
Plateau induction
Light +
hyperpol
2
7
14
50 mV
1 s
-50
-60
-70
0
90
180
position (cm)
15
10
5
0
-5
-4 -2 0 2 4
time from plat. (s)
C
control
10 mV
-4 -2 0 2 4
Time from plat. (s)
D
DG
Plateau induction
Light +
hyperpol
2
8
17
50 mV
1 s
-45
-55
-65
0
90
180
position (cm)
20
10
0
-10
-4 -2 0 2 4
time from plat. (s)
E
DG
10 mV
-4 -2 0 2 4
Time from plat. (s)
F
EC
10 mV
-4 -2 0 2 4
Time from plat. (s)

**Fig. S8. Effect of optogenetic manipulations on BTSP induction.** (**A**) All changes in Vm ramp for individual neurons from CA3 manipulation (blue). (**B)** (from top to bottom) Sketch of protocol for optogenetic silencing; Vm traces for laps before, during and after induction; Vm ramps for laps before (gray) and after (black) plateau induction and difference between Vm ramp after and before plateau ($\Delta$Vm) for single neuron from control group. (**C**) All changes in Vm ramp for individual neurons from control (black). (**D** and **E**) same as in (B and C) but for DG manipulation group (green). (**F**) All changes in Vm ramp for individual neurons from EC manipulation (red).

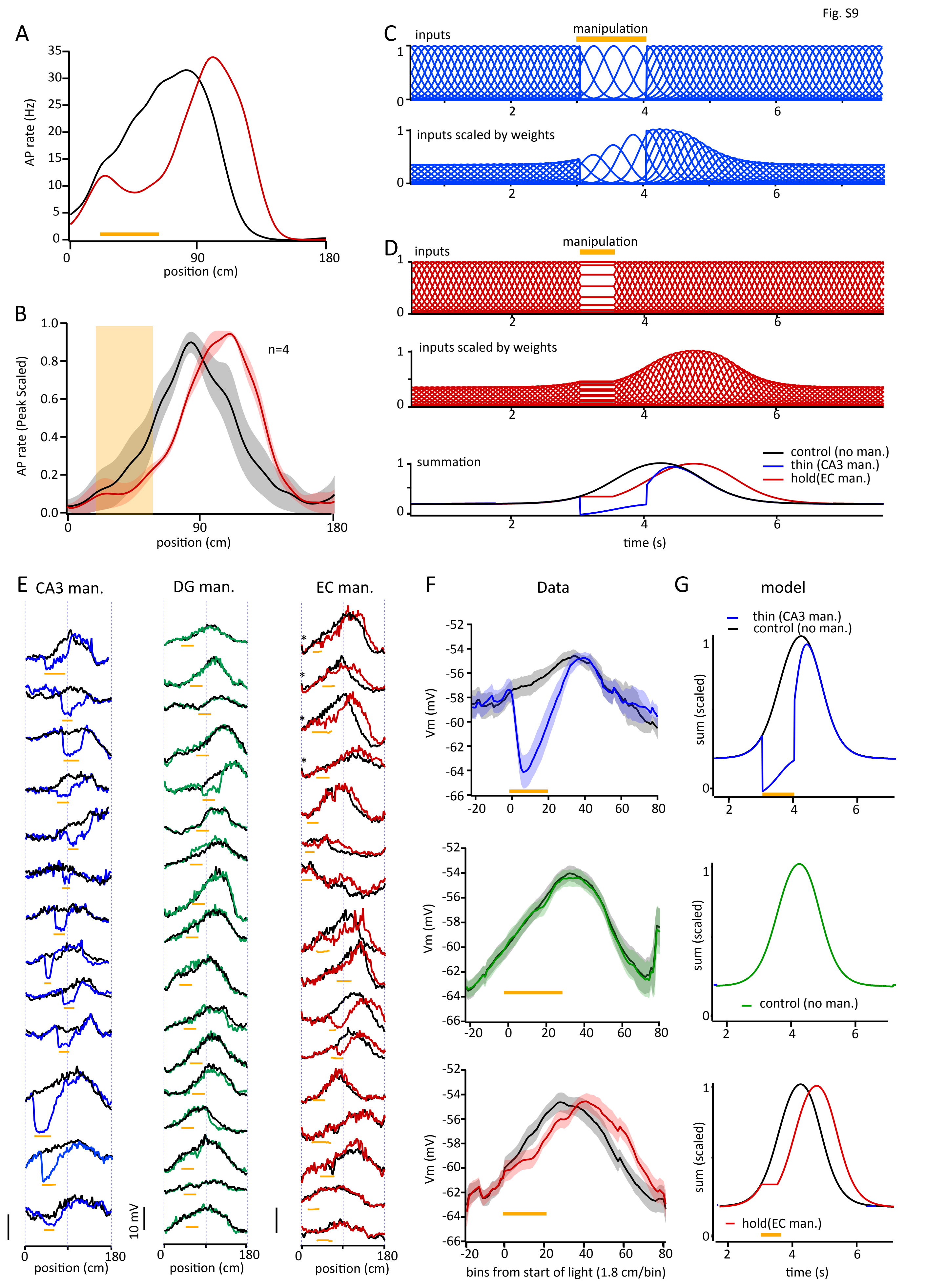

Fig. S9
A
AP rate (Hz)
position (cm)
B
AP rate (Peak Scaled)
n=4
position (cm)
C
inputs
manipulation
inputs scaled by weights
D
inputs
manipulation
inputs scaled by weights
summation
control (no man.)
thin (CA3 man.)
hold(EC man.)
time (s)
E
CA3 man.
DG man.
EC man.
10 mV
position (cm)
F
Data
Vm (mV)
bins from start of light (1.8 cm/bin)
G
model
thin (CA3 man.)
control (no man.)
sum (scaled)
control (no man.)
hold(EC man.)
time (s)

**Fig. S9. Effect of optogenetic manipulations on CA3 PC activity.** (**A**) Mean spatially-binned AP rate for control (black) and test (red) trials for EC manipulation in single neuron. (**B**) Mean spatially-binned AP rate for 4 neurons meeting criteria (peak rate > 10 Hz, peak location near 90 cm and light activation around ~30-60 cm; see * in part (E). (**C**) Signals from model containing control weight distributions but with inputs altered to simulate CA3 manipulation (blue) or (**D**) EC manipulation (red). Simulated Vm ramps shown at bottom. Inputs zeroed only during manipulation period (dashed blue line) or from start of manipulation to end of run (solid blue line). (**E**) All recordings shown where black traces are control conditions (average of pre and post light trials) and colored traces are those during light as labeled. (**F**) Group averages for CA3 (top), DG (middle) and EC (bottom) manipulation data for control (black) and light (blue, green and red for CA3, DG and EC, respectively) trials. Vm ramp are aligned to start of light. (**G**) In the same order top to bottom as in (F) are simulations with the same color code.

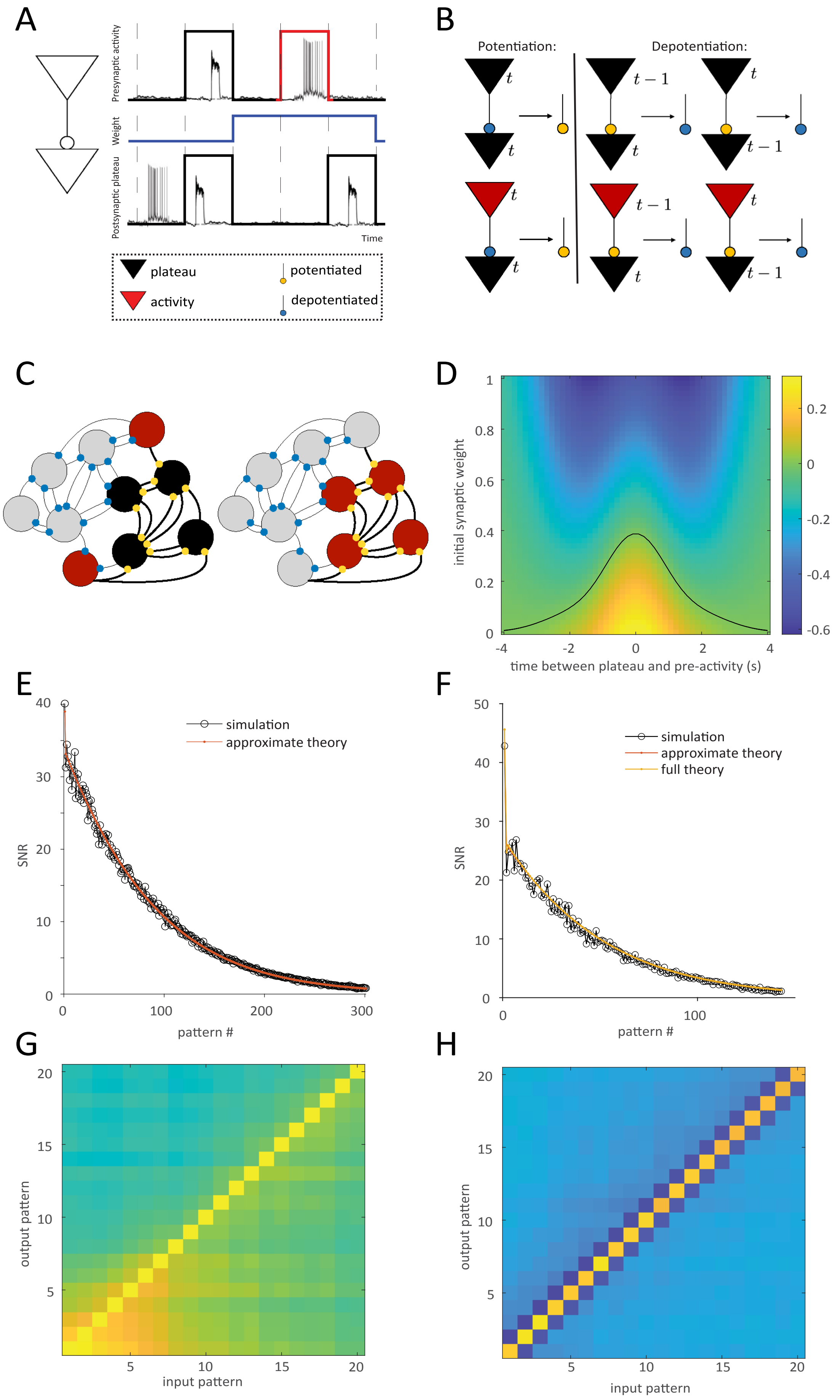
A
Presynaptic activity
Weight
Postsynaptic plateau
Time
plateau
potentiated
activity
depotentiated
B
Potentiation:
Depotentiation:
$t$
$t-1$
C
D
initial synaptic weight
time between plateau and pre-activity (s)
E
SNR
simulation
approximate theory
pattern #
F
SNR
simulation
approximate theory
full theory
pattern #
G
output pattern
input pattern
H
output pattern
input pattern

**Fig. S10: BTSP model.** (**A**) Time discretization and binarization of signals in the binary version of BTSP. Top: Both place field activity (red) and plateaus (black) contribute to the plasticity process in the presynaptic neuron. Bottom: Only plateaus generate plasticity signals in the postsynaptic neuron. Middle: Example synaptic dynamics. An initially depressed synapse (blue, low weight) is potentiated (blue, high weight) in the presence of presynaptic and postsynaptic plateaus in the same time bin and subsequently depressed when presynaptic activity is followed by a postsynaptic plateau. (**B**) Complete binary BTSP rule. Left: A depressed synapse (blue) is potentiated (yellow) in the presence of presynaptic (plateau or activity, top/bottom) and postsynaptic (plateau only) signals at the same time. Right: A potentiated synapse undergoes depression if a postsynaptic signal (plateau or activity) follows (left) or precedes (right) a presynaptic signal. (**C**) Left: Synaptic motifs learned in the network after the presentation of a single pattern. Potentiated synapses (thick edges) form a self-reinforcing assembly among plateau emitting neurons (black circles). The assembly receives feed-forward input from active neurons (red circles) through potentiated synapses. Right: Subsequent transient reactivation of the units will drive persistent activity in the assembly. (**D**) Continuous time and continuous weight version of BTSP (Eq. 6 in Methods). Change in synaptic weight (color coded), as a function of the time interval between single Gaussian pulses for pre-synaptic and post-synaptic signals (x axis) and initial weight (y axis). Black curve denotes combination of pre-post timing and initial weight resulting in no weight change. (**E**) Signal to noise ratio estimated from simulations of independent synapses undergoing BTSP (Eq. 13 in ST, black circles), and from the approximation based on the second leading eigenvalue of the transition matrix from the corresponding Markovian description of BTSP (Eq. 16 in ST, red dots). The age of the patterns increases from left to right, from the most recently stored, to the most remote. (**F**) Same as (C), but with correlated activity across time bins, and including the full theoretical solution (yellow line/dots). (**G**) Confusion matrix (Eq. 12 in ST) for same-time rule in the presence of temporally correlated activity (Eq. 20 in ST) for the last 20 patterns learned by the network. Note how the presentation of any input pattern to the network elicits a signal similar to the few most recently stored patterns (output). (**H**) Same as (C), for BTSP. Parameters for all panels in Methods and Supplementary Information.

sfig11

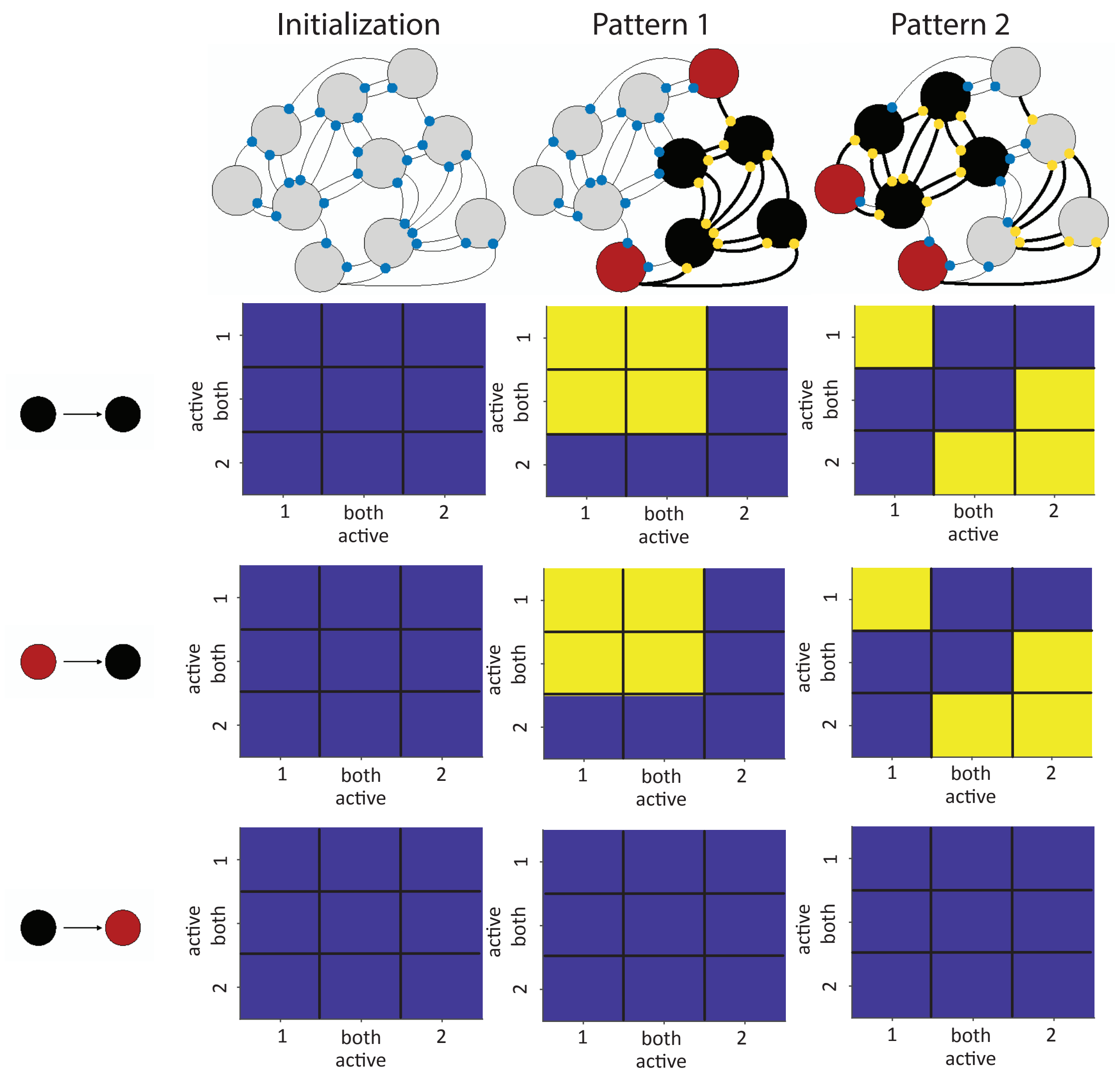

**Fig. S11: Pattern separation of correlated patterns with BTSP.** Evolution of network structure in the presence of BTSP. Top: Left: For simplicity, the network is initialized with depotentiated synapses (thin edges). Middle: Upon presentation of the first pattern (black: plateaus, red: activity), the subset of units with plateaus forms a mutually reinforcing group (thick edges) that receives feed-forward input from active units. Right: Upon presentation of a second pattern, weights among the overlapping plateau and active units across the two patterns are depotentiated, effectively reducing the interference between the first and second pattern (right). Bottom matrices: Weights among units with plateaus are symmetric, forming an assembly of self-reinforcing activity (2$^{nd}$ row). Weights from active units provide feedforward activity to the assembly (3$^{rd}$ row), but do not participate in the assembly (4$^{th}$ row).
